# Memory-bound *k*-mer selection for large and evolutionary diverse reference libraries

**DOI:** 10.1101/2024.02.12.580015

**Authors:** Ali Osman Berk Şapcı, Siavash Mirarab

## Abstract

Using *k*-mers to find sequence matches is increasingly used in many bioinformatic applications, including metagenomic sequence classification. The accuracy of these down-stream applications relies on the density of the reference databases, which, luckily, are rapidly growing. While the increased density provides hope for dramatic improvements in accuracy, scalability is a concern. Reference *k*-mers are kept in the memory during the query time, and saving all *k*-mers of these ever-expanding databases is fast becoming impractical. Several strategies for subsampling have been proposed, including minimizers and finding taxon-specific *k*-mers. However, we contend that these strategies are inadequate, especially when reference sets are taxonomically imbalanced, as are most microbial libraries. In this paper, we explore approaches for selecting a fixed-size subset of *k*-mers present in an ultra-large dataset to include in a library such that the classification of reads suffers the least. Our experiments demonstrate the limitations of existing approaches, especially for novel and poorly sampled groups. We propose a library construction algorithm called KRANK (K-mer RANKer) that combines several components, including a hierarchical selection strategy with adaptive size restrictions and an equitable coverage strategy. We implement KRANK in highly optimized code and combine it with the locality-sensitive-hashing classifier CONSULT-II to build a taxonomic classification and profiling method. On several benchmarks, KRANK *k*-mer selection dramatically reduces memory consumption with minimal loss in classification accuracy. We show in extensive analyses based on CAMI benchmarks that KRANK outperforms *k*-mer-based alternatives in terms of taxonomic profiling and comes close to the best marker-based methods in terms of accuracy.

## INTRODUCTION

The number of genomes available in public repositories has been growing dramatically in recent years, especially due to increased sequencing of microbial and viral species. Of the 322,193 bacterial species deposited on RefSeq since 2000, *>*15% have been added in 2023 alone. A major benefit of having access to all these genomes is to build ultra-large reference libraries with the potential to search new *query* sequences against them. Such libraries can be used to classify reads from a metagenomic sample and to detect contaminants. It has been long appreciated that the accuracy of these downstream applications in metagenomics relies on having access to dense reference libraries as classification accuracy suffers when the query is distant from all reference genomes (Meijenfeldt et al., 2019; Pachiadaki et al., 2019; Rachtman, Balaban, et al., 2020; Liang et al., 2020). Many authors have recently attempted to create ultra-large genomic reference sets (e.g., Parks et al., 2018; McDonald et al., 2023), often used with marker genes (Asnicar et al., 2020; Balaban et al., 2023). Using these datasets with *k*-mers-based methods, however, runs into a mundane but key limitation — the memory needed to utilize these genomes as reference libraries.

Using *k*-mers to build ultra-large libraries and classify reads has been a promising approach, as evident from the success of Kraken 2 (Wood et al., 2019). Kraken 2 and other methods such as CLARK (Ounit and Lonardi, 2016) and CONSULT-II (Ş apcı et al., 2024) extract *k*-mers for some large *k* (e.g., 31) from the reference set, use some strategy to choose a subset of *k*-mers, and build a searchable data structure (e.g., a hash table) of the selected *k*-mers and their taxonomic associations. At the time of the query, the entire data structure is loaded into the memory, *k*-mers from reads are extracted and searched against the data structure, and a read is classified at some taxonomic rank if certain heuristic conditions (specific to each method) are met. It is easy to store millions or even a few billion *k*-mers in memory but modern datasets can include tens or hundreds of billions of *k*-mers. For example, Ş apcı et al. (2024) were able to fit 8 billion 32-mers and their taxonomic labels into ≈140Gb of memory, but these had to be subsampled (arbitrarily) from 20 billion 32-mers available in the moderate-size database of 10,575 genomes produced several years ago by Zhu et al. (2019). Trade-offs of accuracy and scalability imposed by the need to subsample *k*-mers will be increasingly felt by developers of *k*-mer matching methods and those aiming to build ultra-large reference sets.

For the *k*-mer-based methods to benefit from the ever-growing set of available genomes, we need improved methods of subsampling *k*-mers in a memory-bound fashion. Of course, we are not the first to note the need to subsample *k*-mers. Minimization is the standard technique that selects some among adjacent *k*-mers (Zheng et al., 2023; Roberts et al., 2004) and is adopted by most tools. However, we argue that beyond minimization, which focuses on the redundancy of adjacent *k*-mers, we need to consider the evolutionary dimension. This direction can be stated as a computational problem: We are given a taxonomic tree 𝒯 and a set of genomes 𝒢 labeled by the taxonomy. Let 𝒦 be the set all of distinct *k*-mers across all genomes *g* ∈ 𝒢. We seek a subset of 𝒦 with size *M* ≤ |𝒦| such that the reference taxonomy is represented well with the selected *k*-mers. Users can adjust *M* to control the memory budget (see METHODS for details).

The problem statement leaves the meaning of “well-represented” unspecified as many criteria can be considered, and the choice of the criterion forms the basis of each method. A notable attempt in this direction is pioneered by Lee et al. (2011) who propose seeking discriminative *k*-mers specific to a species not found in others, an idea implemented in CLARK (Ounit and Lonardi, 2016). Another obvious approach is to represent all reference genomes at equal levels, or similarly, to sample *k*-mers uniformly at random. The overall goal of this work is to explore these and a set of alternative strategies in the context of taxonomic read classification and taxonomic profiling of metagenomic samples.

Subsampling *k*-mers from ultra-large and heterogeneous databases available these days faces many challenges. On the one hand, sampling of taxonomic groups in public libraries is very imbalanced, which can become a challenge as noted by Nasko et al. (2018). For example, RefSeq includes 35947 *Escherichia* genomes while 1164 genera have only a single genome. In the dataset of Zhu et al. (2019) (WoL-v1 hereafter), despite their attempts to find representative species, the number of genomes sampled per taxon in each taxonomic rank varies three orders of magnitude (Fig. 1A). The true imbalance in evolutionary history (where some clades are much more diverse, abundant, and species-rich than others) in addition to biases in sampling (where some environments are sampled far more than others) are behind such imbalances. Regardless of the cause, a naive subsampling of *k*-mers can easily miss entire branches of the evolutionary tree. The situation is made worse by the fact that for *k* ≫ 20, a large number of *k*-mers are not shared even among genomes of the same species. Examining WoL-v1 dataset, we see that 15% of pairs of genomes from the same species have pairwise distances above 5% (Fig. 1B), which results in 98% of 30-mers being unique to each genome in such pairs (Fig. 1C). Pairs of genomes from the same genus but different species shared more than 10% of their 30-mers in only 1% of cases. Thus, as the number of genomes *N* grows, the number of unique *k*-mers grows close to proportionally to *N*. Moreover, *k*-mers unique to ranks above species will be far outnumbered by those unique to a genome or a species. As a result, optimal *k*-mer selection is even more challenging if one tries to find a unique set of *k*-mers to be used at all ranks (perhaps the reason Ounit and Lonardi (2016) target a particular rank).

**Figure 1:**
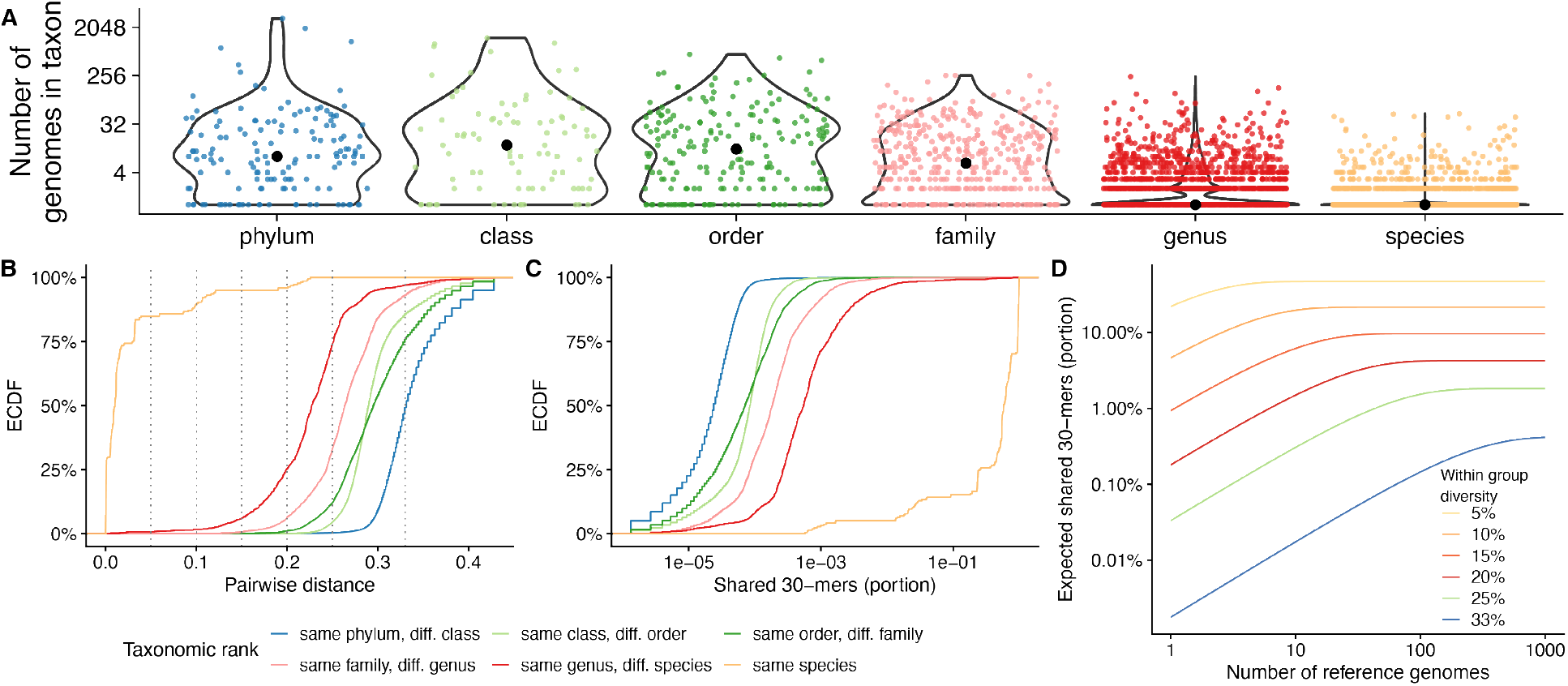
A) Number of genomes in WoL-v1 (Zhu et al., 2019) reference set under each taxonomic node (dots), separated by ranks. B,C) The distribution of Mash (Ondov et al., 2016) estimated genomic distances (B) and Jaccard similarities (C) among 500,000 randomly sampled pairs of genomes that share a taxonomic rank but are different in lower ranks. The empirical cumulative distribution function (ECDF) is shown. D) The theoretical expectation for the number of 30-mers shared between a query and at least one of *N* sampled genomes of a reference set for a group that has within-group diversity 2*d* is shown as (1 − *d*)^*k*^ 1 − (1 − (1 − *d*)^*k*^)^*N*^).

In this paper, we propose a set of strategies for selecting a predefined number of *k*-mers from a large pool of genomes in ways that are conducive to classifying reads. Our goal is to find subsets that do not leave out poorly sampled groups, work across taxonomic ranks, and do not reduce the ability to classify relatively novel reads. We implement and explore these strategies paired with the read classification and taxonomic profiling method CONSULT-II. Our method, *k*-mer RANKer (KRANK), takes a taxonomy and a set of genomes labeled with the taxonomy as input. It selects a subset of *k*-mers from these genomes based on its ranking strategy and a user-defined size constraint. KRANK subsets *k*-mers in a top-down traversal of the reference taxonomy, enabling it to make choices about what *k*-mers to keep locally instead of globally and eliminating the need to analyze all *k*-mers jointly at any point in library construction. Our highly optimized and flexible C++ implementation allowed us to compare several alternative strategies empirically. Our comprehensive results on taxonomic read classification and taxonomic profiling tasks show that KRANK, unlike more naive selection strategies, can produce dramatic reductions in library sizes without substantial loss of accuracy.

## RESULTS

We designed KRANK (detailed in Algorithm 1) to select a predefined number of *k*-mers from a given set of genomes. KRANK designates some taxonomic rank as the leaf rank (species by default). It then traverses the taxonomy 𝒯 in post-order. At leaves, distinct *k*-mers are extracted and encoded in a compact manner (see Algorithmic details of KRANK). Following CONSULT-II, KRANK indexes *k*-mers using *l* locality-sensitive hashing (LSH) tables. These LSH tables ensure two *k*-mers with a small Hamming distance (e.g., less than 3) have a high chance of being indexed to the same row in at least one of the tables (see LSH tables for details). This is achieved by selecting *h < k* random but fixed positions of a *k*-mer as its hash key, and each *k*-mer is stored in the row of the table indexed by the LSH key. As we traverse the tree, on each internal node, we adaptively impose a size constraint on its tables. We combine each row of the hash tables of all the children nodes and remove *k*-mers based on a ranking algorithm until the size constraint is satisfied. At the root, the final *l* tables are restricted to *b* columns per row, and each will contain 2^2*h*^*b k*-mers. When not specified, we use *l* = 2, *k* = 32, *h* = 12, *b* = 16. Given the hash tables at the root, we use the existing CONSULT-II algorithm for taxonomic classification and abundance profiling (see CONSULT-II classification and profiling).

The heart of the algorithm is adaptive size constraints and *k*-mer ranking, both of which we empirically motivate first. These explanatory sets of results use a manageable microbial dataset by Zhu et al. (2019) composed of 10,575 genomes (WoL-v1 hereafter). We chose 756 query genomes in total (676 bacterial and 80 archaeal genomes), of which only 10 are present in the reference set. Measuring the novelty by the minimum distance of a query to any reference genome as estimated by Mash (*d*\*), we bin queries into novelty groups (Fig. S1A). We use 150bp error-prone reads simulated from each genome (66667 reads per query genome) and test methods in terms of their accuracy in classifying individual reads. See Datasets and Evaluation metrics for details. After exploratory results, we move to more formal evaluations of the method for two tasks: taxonomic classification of metagenomic reads and taxonomic profiling of metagenomic samples.

### KRANK: motivating the design

#### Adaptive size constraints

We observed wide differences among taxa, even at the same rank, in terms of sampling density (Fig. 1A) and levels of diversity (Fig. 1B). This heterogeneity creates a major challenge in selecting *k*-mers. If we randomly sample *k*-mers, taxa will contribute proportionally to their sampling in the final subset. Given a total budget *M*, in expectation, a taxon *t* would have 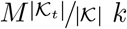-mers in the sampled subset, where 𝒦_*t*_ is the set of *k*-mers of all genomes labeled by taxon *t*, and 𝒦 is the set of all reference *k*-mers. As a result, taxa with lower sampling will have little representation, while highly-sampled groups (e.g., *E. coli*) will dominate. We partially address this challenge by imposing a size constraint on |**H**_*t*_| for each internal node during traversal to avoid having bloated nodes that starve sister taxa. While this balancing can in theory be done at the root, we would need to keep a map from *k*-mers to genomes, which is impractical. Thus, preemptively filtering some number of *k*-mers from some tables helps scalability by reducing the memory usage. We designed a heuristic for adaptive size constraints whereby, each node *t* containing portion *r* (*t*) of the total dataset gets a budget of 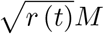 *k*-mers. We define *r* (*t*) as either the portion of total *k*-mers present under *t* (default) or the proportion of taxonomic leaves (i.e., species) under *t* (see Section AdaptiveSizeConstraint for details). We empirically evaluate these two versions of this heuristic against the baseline of enforcing no adaptive constraint, all paired with random *k*-mer subsampling.

Imposing the adaptive size constraints can make a substantial difference for taxonomic groups with low levels of sampling (Fig. 2A). For example for phyla with [0, 50] and (50, 500] reference genomes sampled, both definitions of *r*(*t*) improve the F1 accuracy over enforcing no adaptive constraint for up to 35% and 12%, respectively, with similar improvements obtained at other ranks. Thus, the adaptive size constraints achieve their stated goal of ensuring groups with lower sampling are not left out. This equitable spreading of the *k*-mer budget inevitably results in reduced representation for highly sampled groups. For example, for the few phyla with more than 500 reference genomes, we observe a considerable drop in the performance (e.g., F1 score declines by 14%–20%). Thus, it may appear that better sampling of low-representation groups has to come at the expense of the high-representation groups. However, this tradeoff can be avoided.

**Figure 2:**
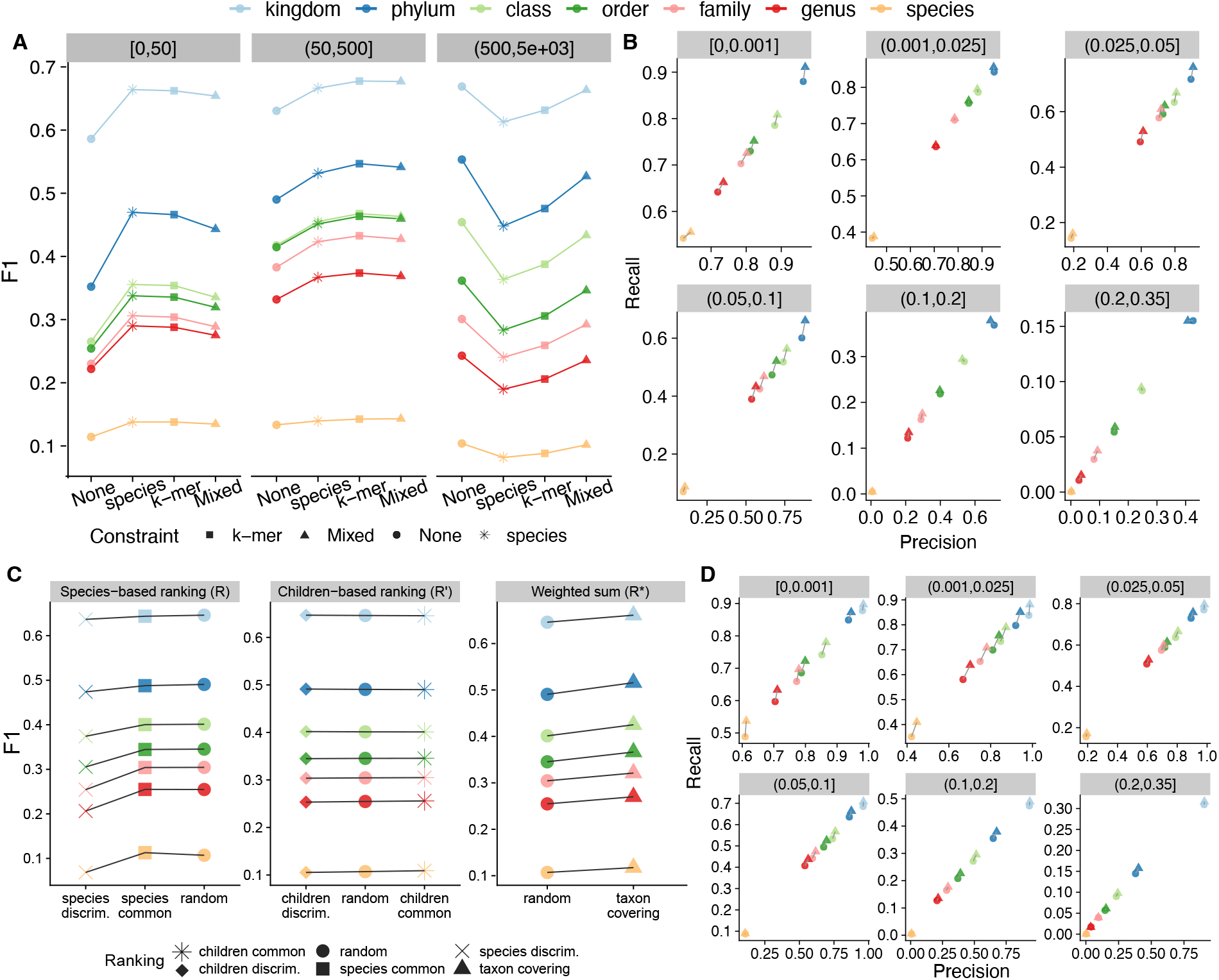
A, B) Comparison of adaptive size-constraint approaches with random *k*-mer selection. None: enforcing no constraint, species: number of species constraint, *k*-mer: total *k*-mer count constraint, Mixed: no constraint for one table and *k*-mer count constraint for the other. A) F1 accuracy, dividing queries (panels) into three bins based on the number of reference genomes in the phylum of the query. B) Precision vs recall of None vs Mixed (default), dividing queries based bins of query novelty (*d*\*). C,D) Evaluation of simple ranking strategies (discriminative, common, random) implemented using species counts R or children counts R^*′*^ and our new weighted-sum R* heuristic (11). See Fig. S2 for an illustration of functions. C) F1 across all queries. D) Precision versus recall of random and weighted-sum heuristic, dividing queries by bins of novelty (*d*\*). In A) and C) *x*-axes are ordered by the average F1 score of all queries. All results use: *w* = 35, *k* = 32, *l* = 2, *h* = 12, and *b* = 16. *d*\* is the distance to the closest reference estimated by Mash. Default KRANK uses mixed adaptive size and R* ranking.

Since we have two different tables (*l* = 2) in the library, we can have a mixed strategy: We enforce the adaptive size constraint for one table but not the other. This mixed strategy is almost as good as the adaptive version for low-representation groups, and almost as good as the no-adaptive-constraint strategy for highly sampled groups (Fig. 2A). Overall, this mixed strategy outperforms having no adaptive constraints, both in terms of precision and recall (Fig. 2B). The improvements are most visible for moderately novel queries (*d*\* ∈ (0.025, 0.1]). Across queries, the mixed strategy improves the recall by 6%–10% and precision by 2%–5% above the genus rank. Thus, KRANK adopts the mixed strategy as the default.

#### Ranking and removing *k*-mers

Faced with the question of which *k*-mers to select given the limited budget, three options immediately present themselves: selecting randomly, selecting *discriminative k*-mers unique to some taxa, or conversely, selecting common *k*-mers present in many taxa. We compared these options and others that we propose paired with the adaptive size constraint approach described above using constraints for both tables (i.e., not mixed). In implementing these strategies, we used two quantities — the number of leaves (i.e., species) under taxon *t* that include a *k*-mer *x* denoted by R(*x, t*), and the number of children of *t* that include the *k*-mer *x* at least once denoted by R^*′*^(*x, t*), as shown in Fig. S2.

#### The case against discriminative *k*-mers

Keeping discriminative *k*-mers found only in specific target species, an approach followed by Ounit, Wanamaker, et al. (2015), would correspond to preferentially filtering out *k*-mers that appear in more than one species (or some other rank), which corresponds to R(*x, t*) *>* 1. Going one step further, we emulate the discriminative *k*-mer strategy by ranking *k*-mers inversely to R(*x, t*) or R^*′*^(*x, t*), thus removing common *k*-mers. Empirically, using either version is worse than simply using random selection (Fig. 2C). The reduction is negligible with children-based R^*′*^(*x, t*) but is quite dramatic when used with species counts R(*x, t*); for example, the declines in F1 are 36% and 19% for species and genus ranks, respectively. Thus, we caution against using discriminative *k*-mers in a memory-bound setting.

The reduced accuracy of selecting unique *k*-mers can be explained by noting that very few *k*-mers are shared even among different species of the same genus. For example, in the WoL-v1 data set (Zhu et al., 2019), a random sample of genomes that are from different species but the same genus share only 0.05% of their 30-mers (Fig. 1C) and have Mash distances that typically exceed 20% (Fig. 1B). With simplifying assumptions, we can approximate the probability that a *k*-mer would be shared between a query genome and at least one of *N* genomes sampled from a taxonomic group, assuming each pair of genomes in this group has distance *d* to parent, as (1−*d*)^*k*^ (1 − (1 − (1 − *d*)^*k*^)^*N*^). Plotting this equation shows that using discriminative *k*-mers leads to very few matches between a query and the reference (Fig. 1D). For instance, in a group with 2*d* = 20% diversity (which is less than most genera; see Fig. 1B), only 0.7% of query 30-mers are expected to be found in at least one among *N* = 5 references genomes and only 4.2% when *N* → ∞ (an infinite number of genomes sampled from that genus). Since most *k*-mers are unique and subsampling is inevitable in a memory-bound setting, removing common *k*-mers makes it difficult to find *k*-mers matches between reads and reference species that have 20% or more distance to them (e.g., when the closest available match is at the genus level). This only gets worse at higher ranks and is not much better within the same species, where the pairwise Mash distance is 5% or more in 15% of cases.

#### Selecting common *k*-mers alone does not help

We can emulate common *k*-mer selection by ranking *k*-mers proportionally to R(*x, t*) or R′(*x, t*). Given the argument against discriminative *k*-mers, common *k*-mers seem appealing because they can represent each taxon using conserved sequences present across different genomes. Empirically, this strategy used with either R(*x, t*) or R′(*x, t*) is better than using discriminative *k*-mers but is no better than random sampling (Fig. 2C). Just like randomly selecting *k*-mers, this strategy focuses the budget on larger and densely sampled taxa. For instance, the WoL-v1 reference set includes 2975 *Pseudomonadota* genomes and only a single genome from many other phyla. While some unevenness is undoubtedly due to genuine differences among the diversity of phyla, some must be due to uneven sampling.

#### Covering all children improves accuracy

Instead of maximizing total coverage, we can attempt to cover all species. We designed a scalable heuristic ranking mechanism to fulfill this goal. We rank *k*-mers by a weighted sum of *R*(*x*,.) values of *t*’s children (see method section FilterByRank and *R*\*(*x, t*) defined in Eq. (11) and illustrated in Fig. S2). The weights are used to de-emphasize children that are highly sampled among surviving *k*-mers. We simply set the weights to be inversely proportional to the *coverage* of each child taxon among *k*-mers of a particular row of the LSH table to ensure that children covered with fewer surviving *k*-mers get covered by the parent. Using this approach improves *k*-mer selection consistently across all taxonomic ranks compared to the random strategy and common *k*-mer strategy (Fig. 2C). Further dividing queries by their novelty, we observe that our weighted sum strategy improves both precision and recall, particularly for less novel queries; recall increases by as much as 15% at the species level in the *d*\* ∈ (0.001, 0.25] bin (Fig. 2D).

#### Comparison to other methods for taxonomic read classification

The default version of KRANK uses both adaptive size constraints (mixed strategy) and *k*-mer ranking based on Eq. (11) and comes in two default modes: high-sensitivity (hs) using 51.2Gb and lightweight (lw) using 12.8Gb. We next compare these two versions of KRANK against CONSULT-II, CLARK, and Kraken 2 on the same WoL-v1 dataset used so far in terms of accuracy in classifying individual reads (Fig. 3).

**Figure 3:**
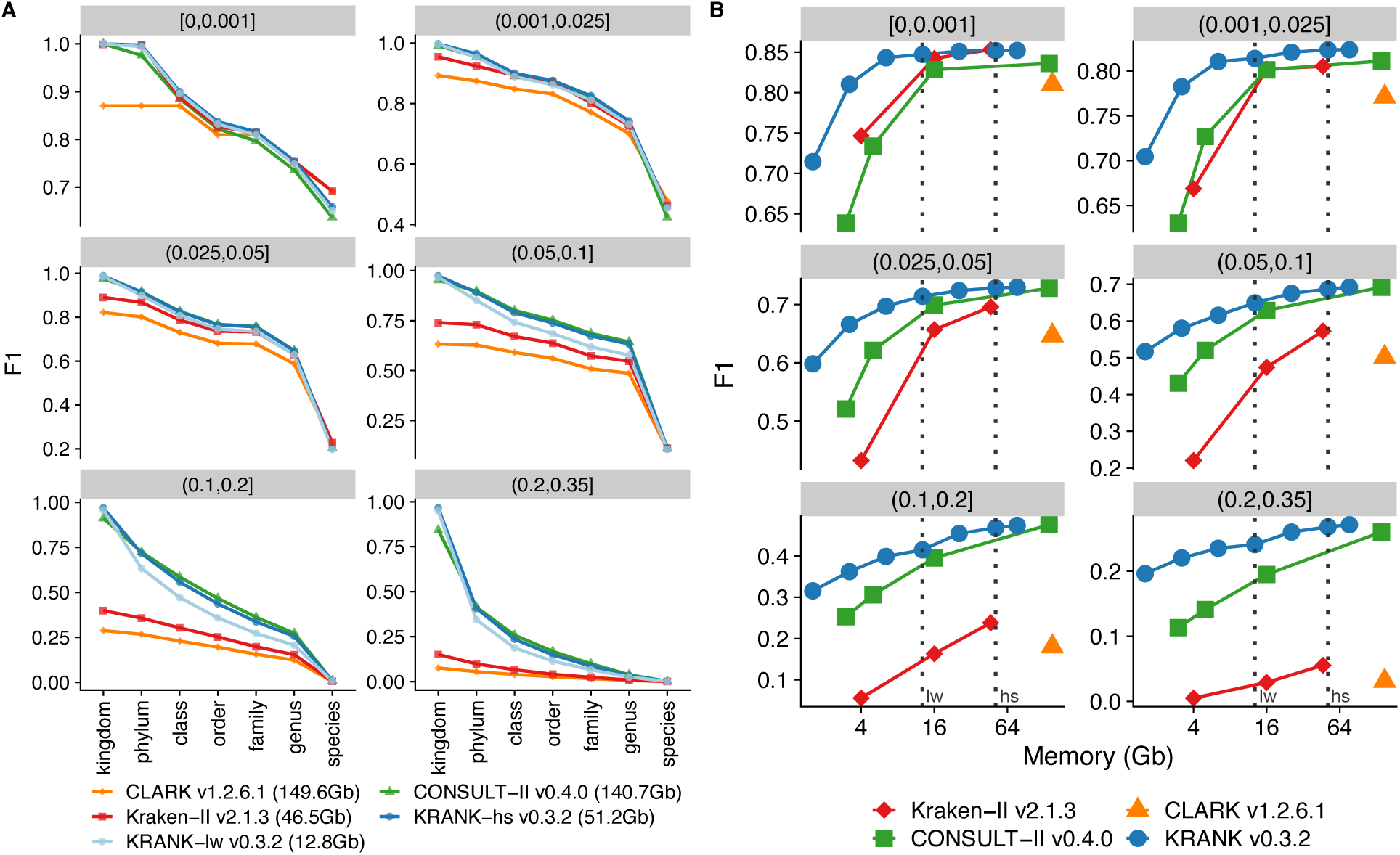
A) Comparison of two default modes of KRANK (lw for lightweight and hs for high-sensitivity) with other methods across ranks and novelty bins on the WoL-v1 dataset with 66667 simulated 150bp reads from each of 756 query genomes. Each panel is a novelty bin, corresponding to the minimum distance to any reference genome (*d*\*), measured by Mash (Ondov et al., 2016); see Fig. S1. F1 scores are averages of all queries in each *d*\* bin. See Fig. S3 for precision and recall. B) F1 accuracy for KRANK, CONSULT-II, and Kraken 2 as we change the memory used in Gb. CLARK is shown as a single data point. We mark memory levels corresponding to the lw and hs modes using dashed lines.

#### Accuracy in default settings

Across different ranks and query novelty levels, KRANK-hs matched or improved on CONSULT-II in terms of F1 accuracy despite using far fewer *k*-mers and about 1*/*3 of the memory (Fig. 3A). KRANK-hs and CONSULT-II often had similar precision (with a slight advantage for CONSULT-II in some cases), but KRANK-hs had better recall in many settings (Fig. S3). Similar to CONSULT-II, KRANK achieved substantially higher recall compared to Kraken 2 and CLARK. Improvement in recall usually comes with little or no sacrifice in precision, which can be controlled with the total vote threshold parameter of CONSULT-II (Ş apcı et al., 2024). KRANK-lw is more accurate than CONSULT-II at the kingdom level and only slightly less accurate elsewhere despite using less than ten times the memory. Accuracy also depended on query novelty (*d*\*). All methods performed similarly both in terms of precisin and recall for queries that were not novel (*d*\* ≤ 0.025), except CLARK which had slightly lower accuracy (Fig. S3). For these less novel queries, both variants of KRANK were indistinguishable from CONSULT-II. For more novel queries (*d*\* *>* 0.025), accuracy dropped for all methods and substantial differences between them emerged. For example, for the *d*\* ∈ (0.05, 0.1] bin, KRANK-lw outperformed Kraken 2 and CLARK above the species rank, achieving 16% and 35% higher F1 scores, respectively, at the phylum level. Unlike KRANK-hs, KRANK-lw had slightly lower accuracy than CONSULT-II at some ranks (e.g., 8%–9% at the order rank) for *d*\* ∈ (0.025, 0.1] and moderately lower accuracy (e.g., 12% at the phylum rank) for *d*\* *>* 0.1. Nevertheless, KRANK-lw out-performs Kraken 2 (e.g., 77% and 250% higher F1 at the phylum rank for *d*\* ∈ (0.1, 0.2] and (0.2, 0.35], respectively).

Both KRANK-hs and KRANK-lw are slightly better than CONSULT-II in terms of kingdom-level classification for novel genomes. This improvement can be due to the better representation of archaea with careful *k*-mer selection. Note that our query set is highly imbalanced (e.g., 90% bacteria and 33% *Pseudomonadota*). Thus, simply assigning all reads to the largest group (in terms of the number of query genomes) at each rank would still perform well (Fig. S4). As expected, this null model performs well at the kingdom level (0.94 F1 score). However, KRANK (both lw and hs versions) is the only method that improves on the null model (0.96 and 0.97 average F1 scores, respectively) and assigns 83% of archaeal reads to archaea. At lower ranks, all tools, except for CLARK at the phylum rank, do substantially better than the null model.

#### Adjusting memory usage

We next explored three to seven levels of consumed memory for KRANK, CONSULT-II, and Kraken 2 to compare their accuracy given similar levels of memory, ranging from 1.6Gb to 140Gb (Fig. 3B). KRANK clearly and dramatically outperformed both alternatives in terms of average F1 score when memory is controlled. At lower memory levels, CONSULT-II had much lower F1 scores than KRANK, and Kraken 2 had lower F1 than CONSULT-II for more novel queries. For instance, given ≤4Gb, KRANK was 26% more accurate than CONSULT-II and 8% more accurate than Kraken 2 for the least novel bin. For the moderately novel queries (*d*\* ∈ (0.05, 0.1]), KRANK with 3.2Gb outperformed Kraken 2 with 46.5Gb. KRANK with 1.6Gb outperformed CLARK (149.6Gb) and Kraken 2 (46.5Gb) for novel queries (*d*\* *>* 0.1), giving 160% and 89% higher F1 scores, respectively. Note that CONSULT-II and KRANK store the same number of *k*-mers (268M in a single table) when memory usages are 5Gb and 3.2Gb, respectively. KRANK remained more accurate than CONSULT-II when we compared the method for varying numbers *k*-mers kept in the library (Figs. S5 and S6). Since the only difference between KRANK and CONSULT-II is the selected *k*-mers, these results demonstrate the effectiveness of the selection algorithm.

#### Resource usages

Despite performing more complex computations, KRANK built libraries using considerably less computational resources due to its efficient implementation, compared to CONSULT-II (Table 1). While the total time needed was also less, the main gain was KRANK’s distributed memory implementation. KRANK splits the LSH table into batches, enabling it to divide the workload into independent jobs. For instance, on WoL-v1 dataset, KRANK-hs was built using 512 batches per table, each of which took 8 minutes on average with 4 threads. KRANK-lw used 256 batches and built each batch in 5 minutes on average. Together with initial preprocessing of reference genomes, processing 32 batches in parallel (e.g., on different cluster nodes) needed 4.5 hours for high-sensitivity mode to construct a library with two tables. Using this configuration, the peak memory usage for each batch was less than 32Gb. Although the running times are not directly comparable because of the difference in parallelism, KRANK needed significantly shorter times than CONSULT-II and CLARK to build its libraries.

**Table 1:** Computational resources needed for a database built on 10,470 reference genomes and 756 queries (50M short reads). Measurements for both queries and library building were performed on a machine with 2.2 GHz AMD EPYC 7742 processors. Reported library building times are for 256 and 512 batches, respectively for KRANK-lw and KRANK-hs, each with 4 threads, distributed across 32 cluster nodes and run in parallel.

|  | Running time |  | Peak Memory |  |
| --- | --- | --- | --- | --- |
|  | Query | Library | Query | Library |
| Kraken 2 | 88 sec, 96 threads | 3.5 hr, 128 threads | 47Gb | 50Gb |
| CLARK | 676 sec, 96 threads | 13.5 hr, 128 threads | 150Gb | 370Gb |
| CONSULT-II | 2191 sec, 96 threads | 19 hr, 128 threads | 141Gb | 157Gb |
| KRANK-hs | 1167 sec, 96 threads | 4.5 hr, 32×4 threads | 51Gb | 32Gb per batch |
| KRANK-lw | 787 sec, 96 threads | 2.5 hr, 32×4 threads | 13Gb | 8Gb per batch |

At the query time, compared to CONSULT-II, KRANK does not introduce any extra computation but its smaller libraries resulted in reduced running times. On this dataset, KRANK-hs was more than twice as fast as CONSULT-II (Table 1). These improvements are due to better cache performance and shorter library loading times. Note, however, that Kraken 2 was around 13*×* faster than KRANK-hs and scaled considerably better as the number of queried short reads increased (Fig. S7). This difference may be due to the fact that KRANK needs to compute HD for up to 2*b* = 32 reference *k*-mers for each query *k*-mer whereas Kraken 2 only computes presence/absence.

### Abundance profiling: CAMI challenges

#### CAMI-I high complexity dataset

We next compared KRANK-hs against three alternatives in terms of taxonomic profiling accuracy on the Critical Assessment of Metagenome Interpretation (CAMI-I) (Sczyrba et al., 2017) benchmarking challenge, focusing on the high complexity subset (can be found at http://gigadb.org/dataset/100344). We compared against CONSULT-II, CLARK, and Bracken (a Bayesian extension of Kraken 2) using standard metrics promoted by CAMI-I (see Datasets and Evaluation metrics). All methods were run with the custom libraries built using the same WoL-v1 dataset used in the previous analysis.

Abundance profiles estimated by KRANK-hs and CONSULT-II are the most similar to the true profile in terms of Bray-Curtis dissimilarity (Fig. 4A). At the species level, all methods have high errors and are comparable, with a slight advantage for KRANK-hs. As we move up the ranks, the error quickly drops for all methods, but KRANK-hs and CONSULT-II outperform others. Overall, KRANK-hs performs similarly to CONSULT-II despite the much-reduced memory. KRANK performs considerably better than both Bracken and CLARK. All methods tend to underestimate Shannon’s equitability, which is a measure of the variety and distribution of taxa present in a sample. At the phylum rank, Bracken has closer values to the gold standard, while CONSULT-II and KRANK become better at the family rank or lower (Fig. 4B). Overall, KRANK-hs is slightly less accurate than CONSULT-II in terms of this metric across ranks. Note that correcting for genome size improves the accuracy of both CONSULT-II and KRANK, but interestingly KRANK benefits slightly more in terms of the Bray-Curtis metric (Fig. S8A). Since the largest three phyla in our reference set constitute 80%–85% of each sample, this dataset presents a case where the KRANK strategy of covering low-sampled groups is not needed and sampling of highly covered phyla by CONSULT-II is sufficient.

**Figure 4:**
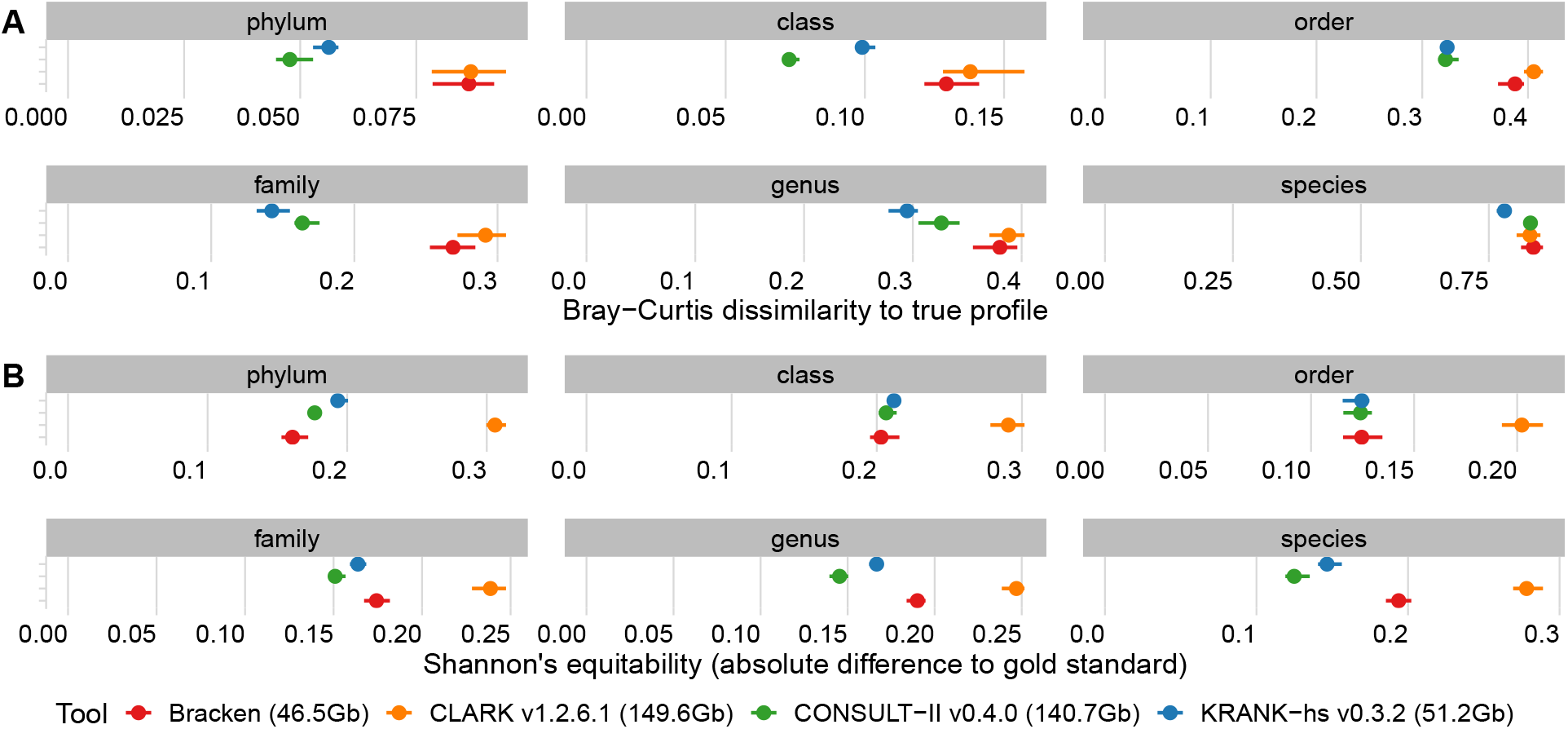
Taxonomic profiling on CAM-I challenge. A) The Bray-Curtis metric measures the dissimilarity between the estimated and true profiles. B) The difference between Shannon’s equitability of the estimated profile and of the gold standard, measuring how well each tool reflects the alpha diversity (i.e., the evenness of taxon abundances). Reported metrics are based on the profile estimates of the high-sensitivity (hs) setting of KRANK. We show the mean and minimum maximum across five samples.

#### CAMI-II marine and strain-madness datasets

To compare KRANK to a larger set of methods, we built a library based on 72,766 genomes available on RefSeq as of 2019/01/08. Since this same dataset had been provided to all methods tested in the CAMI-II challenge (Meyer, Fritz, et al., 2022), we were able to compare KRANK with 12 alternative methods from the original paper, in addition to CONSULT-II, which was also applied to this dataset (see Datasets). These methods include both marker-based and *k*-mer-based methods, which use reads from the entire genome as opposed to relying on predefined markers.

Taxonomic profiles of KRANK-hs were among the most accurate on the CAMI-II challenge (Fig. 5). On the strain-madness dataset, in which only 2 phyla constitute 90% of the relative abundance, our method was a close second to the best method, which was the marker-based method MetaPhlAn (Figs. 5 and S9A). Averaged across all samples, the UniFrac score of KRANK-hs was only 2% less than MetaPhlAn and 6% above the third-best method, CONSULT-II. In species, class, and phylum ranks, MetaPhlAn had slightly better L1 than KRANK-hs, while in the family rank, KRANK-hs was slightly better (Fig. 5). Despite using the same profiling algorithm as CONSULT-II and 1*/*3 of its memory, KRANK-hs also had better L1 metrics than CONSULT-II across all ranks on this dataset. Other *k*-mer-based methods such as Bracken had far lower accuracy in terms of both metrics.

**Figure 5:**
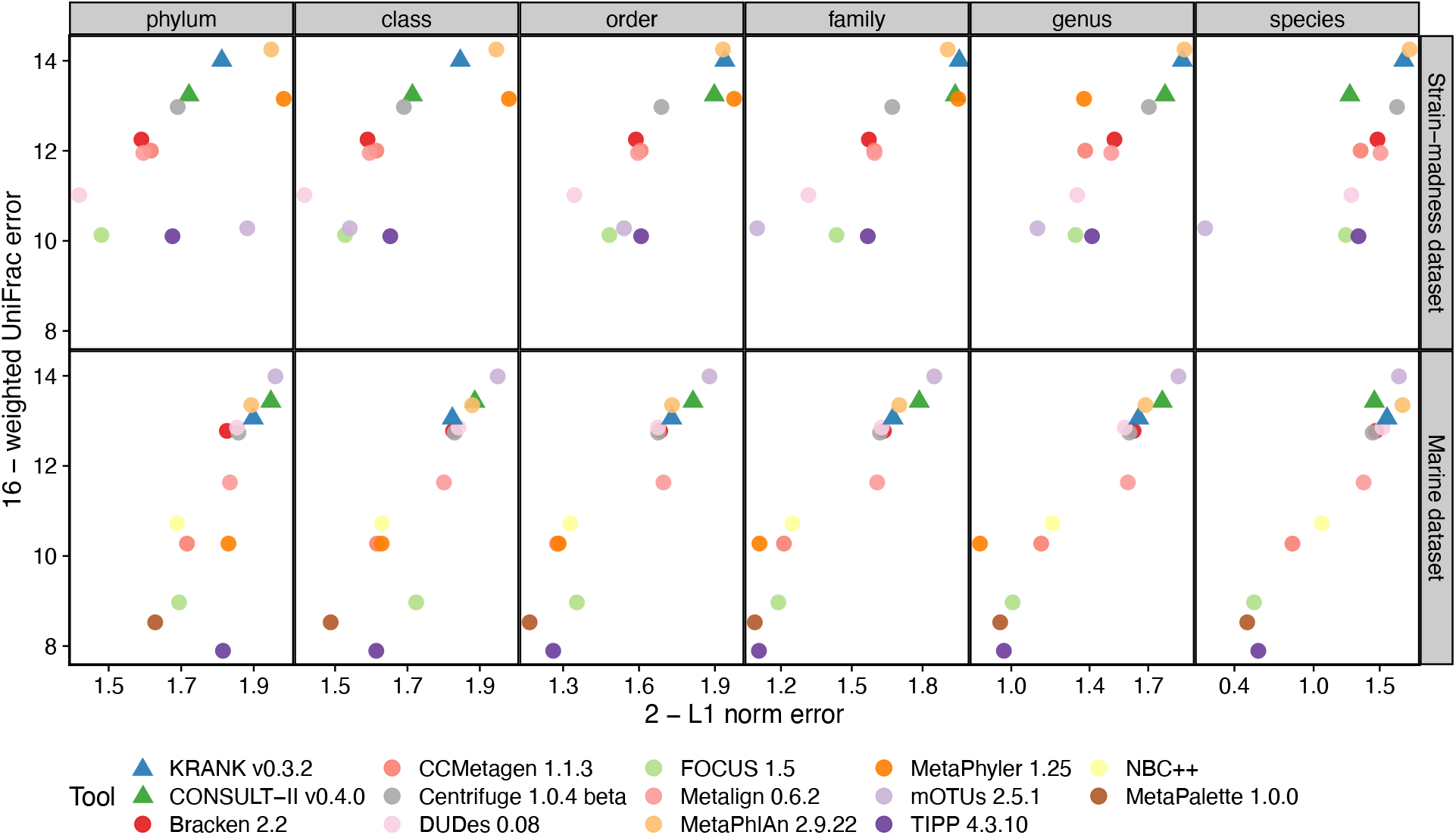
Comparing KRANK (high-sensitivity setting using 51.2Gb) with other participants in CAMI-II benchmarking challenge (Meyer, Fritz, et al., 2022). As in the original manuscript, we show the upper bound of L1 norm (2) minus actual L1 norm versus the upper bound of weighted UniFrac error (16) minus actual weighted UniFrac error. Each data point stands for the average of 100 strain-madness samples or 10 marine samples. Metrics were computed using OPAL with default settings and the -n option. The UniFrac error is the total amount of predicted abundances that needs to be moved along the edges of the taxonomic tree to make them overlap with the true abundance profile. The L1 error simply measures the accuracy of reconstructing the relative abundance profile at a fixed rank.

On the marine dataset, which includes both archaeal and bacterial genomes, and consists of 65%– 70% *Proteobacteria*, the pattern slightly changed (Fig. 5). Here, CONSULT-II was slightly more accurate than KRANK-hs, with 3% better UniFrac and 3% better L1, averaged over all ranks and samples. Only at the species level KRANK-hs was more accurate than CONSULT-II (6% in terms of L1). Overall, KRANK-hs was ranked fourth, following mOTU, CONSULT-II, and MetaPhlAn (Fig. S9). Note, however, that the best method, mOTU, performed poorly on the strain-madness dataset. KRANK-hs was the second best *k*-mer-based method after CONSULT-II but used only 1*/*3 of its memory. The third leading *k*-mer-based method, Bracken, was less accurate than CONSULT-II and KRANK-hs in both metrics, but with a smaller margin than the strain-madness dataset.

To summarize, KRANK-hs was the best *k*-mer-based method and the second best method overall (Fig. S9B). Averaged over both datasets, the marker-based method MetaPhlAn had the best accuracy, followed closely by KRANK-hs, and then CONSULT-II. Note, however, that we focused on the two main CAMI-II metrics that accounted for abundance values. Judged by the presence/absence of taxa (i.e., ignoring abundance), KRANK-hs had better purity than CONSULT-II and Bracken and about the same level of completeness (Fig. S10). However, these abundance-agnostic metrics were not the strength of *k*-mer-based methods; across both datasets, KRANK-hs, CONSULT-II, and Bracken were among the best in terms of completeness but not purity.

## DISCUSSION

We introduced KRANK for selecting a memory-bound and representative subset of *k*-mers from a large genome collection for the purpose of taxonomic classification and abundance profiling. We used a tree-based *k*-mer filtering algorithm that decides which *k*-mers to remove as it moves up a taxonomic tree. This bottom-up strategy makes the library construction more memory efficient as *k*-mers are analyzed in smaller chunks at each node. In addition, we use an adaptive size constraint mechanism to leave some filtering decisions to higher taxonomic levels. These decisions are made based on the prevalence of *k*-mers among children of or leaves under each node.

Ultimately, both our adaptive size constraints and our ranking strategies are heuristics, with no theoretical guarantees, but designed based on careful experiments involving novel queries and imbalanced sampling. Due to the empirical nature of the method, the exploration of alternatives will be an ongoing topic of research, one that is helped by KRANK’s software design. In particular, our approach still leaves room for ties during ranking, and smarter ways to break such ties may help accuracy. Alternatively, more theoretically justified approaches should be explored.

One obstacle to building a more theoretical approach is to define an objective function for *k*-mer selection. Subtleties of imbalanced and biased sampling in reference databases make the definition of the desired optimization problem tricky. Should we attempt to combat the sampling bias by selecting *k*-mers more proportionally from under-sampled groups? If so, how can we know and model the bias? Should we attempt to consider likely profiles of queries? If so, under what model? These questions have no easy answers. Nevertheless, defining mathematically rigorous but nuanced objectives for *k*-mer selection could be a fruitful avenue of future research.

Our approach focused on *k*-mer similarity along the evolutionary dimension but completely ignored *k*-mer overlap. Selecting *k*-mers based on overlap is well covered by minimizers, which we use as a preprocessing step. However, our ranking algorithm is unaware of overlap and minimizers are unaware of the taxonomic relationships. Integrating the evolutionary-aware strategies of KRANK with location-based minimizers poses an interesting algorithmic challenge and could further improve the method. However, designing such a combination is not trivial given our hashing strategy and needs further algorithmic developments.

Despite all the improvements compared to CONSULT-II, the library construction of KRANK is expensive in terms of CPU hours spent. Nevertheless, the creation of the reference library is a one-time operation, amortized over many subsequent uses. Furthermore, computation is getting cheaper rapidly, and the total resources we used (e.g., less than 550 CPU hours for the WoL-v1 library) are far from prohibitive. Crucially, the batching strategy enables us to build the library over a cluster (i.e., distributed memory), with each job using as little as 8Gb and many running in parallel in the order of minutes (Table 1). Moreover, note that the library building time is a function of both the number of genomes and the size of the taxonomy, which grows less rapidly. For example, while the CAMI-II library had 7*×* more genomes than WoL-v1, we needed only 3*×* times, since the taxonomy had 1.8*×* as many nodes, and 2*×* as many species as WoL-v1. Our analyses of 72,766 genomes included in the CAMI-II case demonstrate the scalability of the method to scales commensurate with modern-day libraries.

As datasets become even larger, KRANK parameters may need to adjust. Many ultra-large reference databases are available now, including GTDB (Parks et al., 2018) with 402,709 genomes, and WoL-v2 (Balaban et al., 2023), with a phylogeny built from 199,330 genomes. Moreover, RefSeq currently includes millions of genomes. The fast increase is primarily due to the sequencing of many more genomes from key species. As the number of genomes from each species increases, our choice in this paper to treat the species rank as the leaves of the taxonomy becomes less scalable. However, KRANK can easily use a more refined group (e.g., subspecies) as the leaf, which will trade off slightly longer running times for better memory efficiency. It could also enable sub-species-level classification, a topic that should be explored in the future. Another concern in handling ever-growing datasets is the need for *de novo* construction of the database as new genomes become available. Future research should explore ways to add new genomes into existing libraries, without repeating all the steps.

## METHODS

As a *k*-mer selection method, KRANK can, in principle, be paired with any *k*-mer-based classification or profiling methods. However, in this paper, we use KRANK solely with CONSULT-II (Ş apcı et al., 2024); *k*-mer matches found in libraries built by KRANK are given as input to CONSULT-II v0.5.0 for classification and profiling. Moreover, our *k*-mer selection algorithm adopts a similar LSH data structure to CONSULT-II. Thus, we start by describing the CONSULT-II hash tables and the CONSULT-II classification and profiling methods before presenting the details of KRANK.

### LSH tables

CONSULT(-II) was designed to answer the following question: Are there any *k*-mers in a given reference set with Hamming distances (HD) less than some threshold *p* to a given query *k*-mer? The core idea is to find low HD matches efficiently by partitioning reference *k*-mers to constant-sized subsets (i.e., rows of the hash table with size *b*) using the bit-sampling LSH method (Har-Peled et al., 2012). In this method, we select *h* random but fixed positions of a *k*-mer as its hash key. Each LSH table is a simple 2^2*h*^ *× b* array, and *l* such hash tables constitute the data structure (default: *l* = 2, *h*=14, and *b*=16 for KRANK).

Each query is compared against all *l × b k*-mers with the same hash key. We return the closest *k*-mer match with distance ≤ *p* if any exists. Since we explicitly compute the HD of the query to these reference *k*-mers, there are no false positive matches but false negatives are possible. Using the bit sampling scheme (Har-Peled et al., 2012), one can guarantee that two *k*-mers at HD=*d* have the same hash with probability (1 − *d/k*)^*h*^ assuming independence of positions. Thus, given two *k*-mers, the probability that *at least one* of *l* LSH keys is the same for both *k*-mers is given by

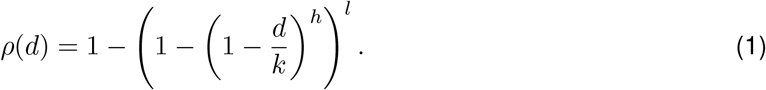

For small enough *p* (e.g., *p*=4), the probability *ρ*(*d*) drops quickly when *d* ≫ *p* and it is sufficiently high if *d* ≤ *p*. Furthermore, since classification is done at the read level, there will be *L* − *k* + 1 chances for a *k*-mer match for a read of length *L*. For such a read, the expected number of matching *k*-mers is (*L* − *k* + 1)*ρ*(*d*), which can be large enough for several realistic choices of *l* and *h* (Fig. S11). For instance, with *k*=30, *h*=14 and *l*=2 (KRANK-hs settings), for a 150bp read at 15% distance from the closest reference genome, we would still expect 23.5 *k*-mer matches *if all of those reference k-mers are in the database*. If only 20% of the *k*-mers from the reference genome are in the library, we still expect more than 5 *k*-mer matches, which can be sufficient for classification. In practice, the number of matches will depend on the budget *M*, and will necessarily decrease as we reduce *M*.

CONSULT-II extends LSH tables further by adding a taxonomic label for each *k*-mer in the table. This label is the lowest common ancestor (LCA) of a *subset* of species that have the corresponding *k*-mer (hence the name *soft* -LCA). This subset is determined with a probabilistic procedure that performs a Bernoulli trial for each genome and includes its species in the subset if the trial is successful. The success probability is different for each *k*-mer *x*_*i*_ and is a function of the number of genomes in which the *k*-mer appears 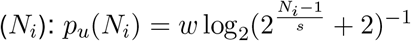, where *w* and *s* are hyperparameters.

### CONSULT-II classification and profiling

Both read classification and profiling algorithms of CONSULT-II are based on voting. Given a query read *R*, CONSULT-II lets each *k*-mer *x* ∈ *ℛ* vote for the soft-LCA of the reference *k*-mer it matches. The value of the vote decreases as the HD between the query *k*-mer *x* and its best match increases. Let the best match be at distance *d* to a *k*-mer with soft-LCA at taxon *t*; then, *x* votes 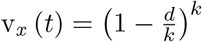 units for *t*. To take hierarchical relations in the taxonomic tree into account, CONSULT-II aggregates votes by recursively summing them in a bottom-up manner. Let child(*t*) be the set of children of the taxon *t*. Then, the total vote of taxon *t* for read ℛ is:

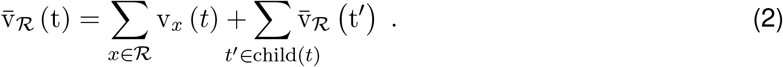

For read classification, CONSULT-II simply determines a majority vote threshold *τ*, corresponding to half of the total vote value at the root of the tree. It then assigns each read to the group at the lowest rank whose total vote value strictly exceeds *τ*. Note that this choice is guaranteed to be unique.

To derive an abundance profile, CONSULT-II first normalizes the total vote values of a read ℛ_*i*_ per each taxonomic rank using

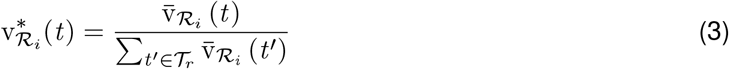

where 𝒯_*r*_ is the set of all taxa at rank *r* (e.g., all phyla). Next, it gathers normalized total vote values of all *n* reads ℛ_1_, …, ℛ_*n*_ in a sample, and normalizes again to obtain the final profile. Let 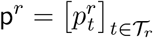 denote the relative read abundance profile at rank *r*, summing up to 1. CONSULT-II (since sets the relative abundance of taxon *t* to:

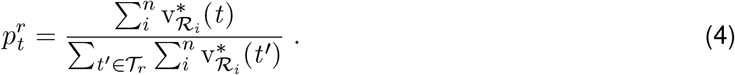

Note that 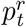 estimates the ratio of *reads* belonging to the taxon *t*, but we are often interested in the relative abundances of *cells* (denoted by 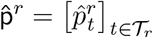. Thus, CONSULT-II (since v0.4.0) incorporates genome sizes by simply correcting 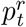 values as follows:

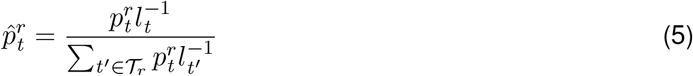

where *l*_*t*_ is the average genome length of all references in taxon *t*. See Fig. S8A for a comparison between v0.3.0 corresponding to Eq. (4) and v0.4.0 implementing Eq. (5) on the CAMI-I dataset. As an option, CONSULT-II can replace the denominator of Eq. (4) with the total vote at the root of the taxonomic tree to quantify the abundance of unclassified taxa. This can be viewed as propagating vote values down from a parent to an artificial “other” lineage which continues until the species rank.

As opposed to classification, CONSULT-II’s profiling (v0.4.0) algorithm does not require a majority vote, and hence, a read can contribute to abundance values of multiple taxa at a fixed rank, even when there exist only a few high HD matches. Furthermore, normalized values can accumulate over billions of reads in a sample, and can result in erroneous profiles (potentially explaining relatively high errors of CONSULT-II at species rank in CAMI-II strain-madness dataset 4). In v0.5.0 (first introduced in the present manuscript), CONSULT-II introduces a new profiling algorithm, which uses a very similar principle to its classification algorithm. Keeping the genome size correction (Eq. (5)) the same, v0.5.0 modifies Eq. (4) as follows:

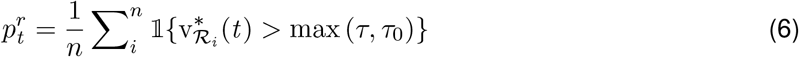

where *τ* is the total vote value of the taxonomic tree and *τ*_0_ is a hyperparameter hard threshold (default: 0.03). In words, instead of each *k*-mer of a read voting for a different taxon, each read votes collectively; the read votes for the same taxon where CONSULT-II would classify it if its total vote is high enough (i.e., *> τ*_0_). Reads with total votes below *τ*_0_ and few or ambiguous matches are considered unclassified. Thus, the resulting abundance profile does not have to sum to 1. Using v0.5.0 instead of v0.4.0 with KRANK’s library resulted in improvements in terms of Bray-Curtis dissimilarity especially at lower ranks (Fig. S8). Thus, we report profiling results using v0.5.0 for KRANK in both CAMI challenges.

#### Algorithm 1

KRANK algorithm. Functions are discussed in the text.

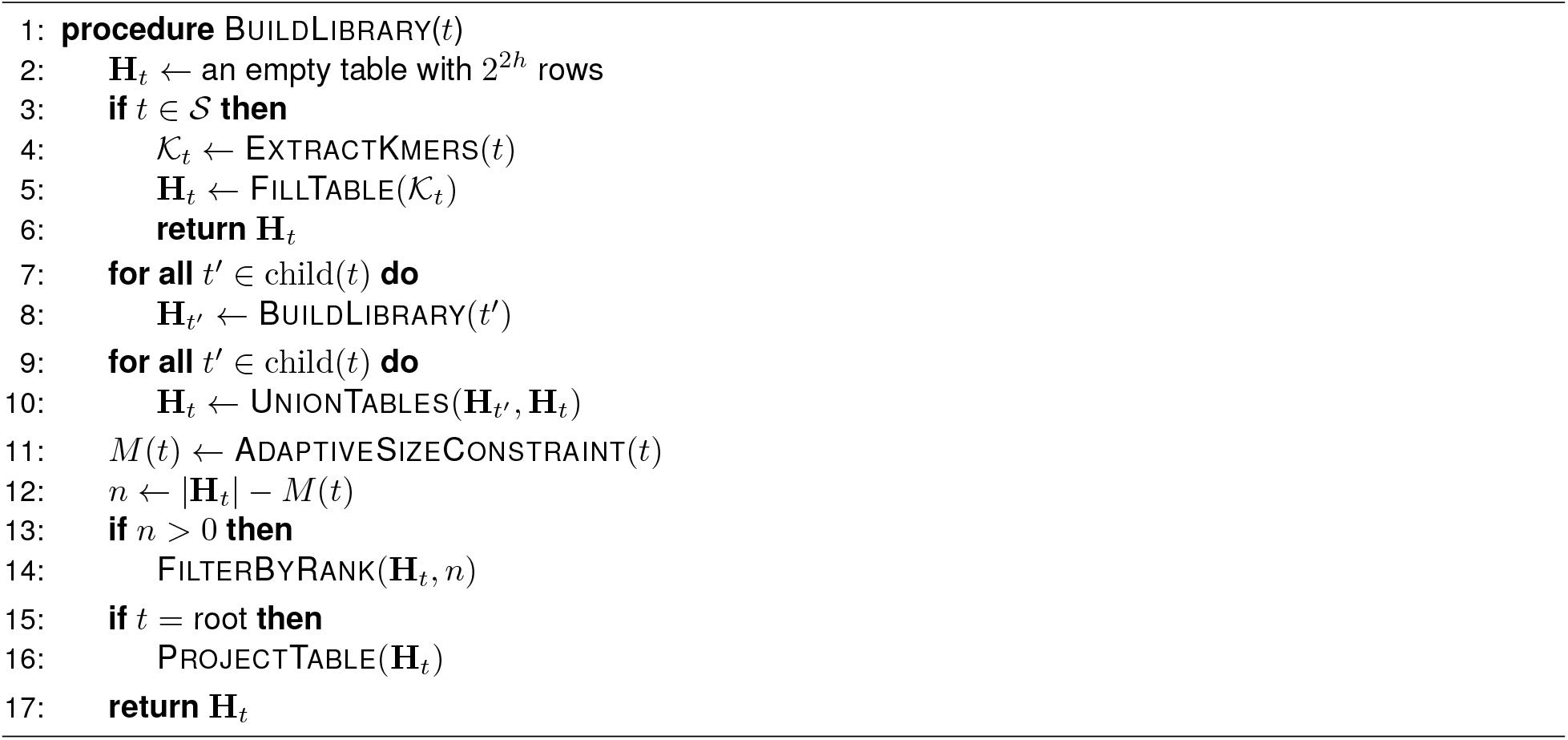

### Algorithmic details of KRANK

We use 𝒯 to denote the set of all nodes of the taxonomy when clear by context. We let 𝒮 ⊆ 𝒯 be the set of species, and for *t* ∈ 𝒯, we let 𝒮_*t*_ ⊂ 𝒮 be the set of species under node *t*. We can choose the leaves of the tree 𝒯 to correspond to any desired rank, but by default, we use species (𝒮) as the lowest rank; this can be easily changed to any other rank. Let 𝒦_*t*_ denote the set of *k*-mers of all genomes labeled by a taxon *t* ∈ 𝒯. Notice that, 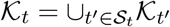.

Recall that during the tree traversal of Algorithm 1, for each node *t* ∈ 𝒯, three key operations are performed to build a new hash table denoted by **H**_*t*_:

1. UnionTables: takes the union of hash tables of its children, **H**_*t*′_, *t*′ ∈ child(*t*),
2. AdaptiveSizeConstraint: computes a size constraint for **H**_*t*_,
3. FilterByRank: ranks and removes *k*-mers from the union until it fits the size constraint.

At the root, the library is flattened to a 1D array (ProjectTable). We first describe our two main heuristics and then discuss details of other named functions in Algorithm 1.

#### AdaptiveSizeConstraint

Let *M* (*t*) ≤ *M* be the upper bound (i.e., the size constraint) for the number of *k*-mers kept for each table at each node (i.e., AdaptiveSizeConstraint in Algorithm 1). Let *r* : 𝒯 → (0, 1] assign a portion of the full budget *M* to each node *t*; any function *r* is valid as long as it takes the value 1 at the root (i.e., *r*(root) = 1) and increases as we move up the tree (i.e., *r*(*t*) ≥ *r*(*t*′) if *t*′ ∈ child(*t*)). We evaluated two options for *r* (*t*):

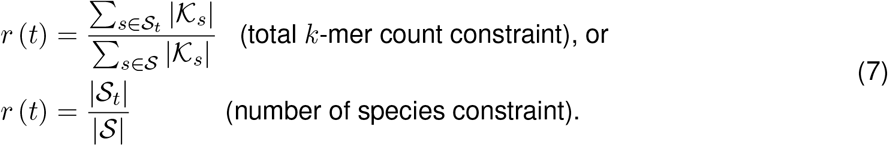

The two options for *r* (*t*) would be identical if all species have equal-sized genomes and each species is sampled the same number of times (𝒦_*s*_ can be the union over multiple genomes). Their main difference is that the first option allows highly sampled species (or very large genomes) to contribute more, while the latter does not. With either definition, we then set

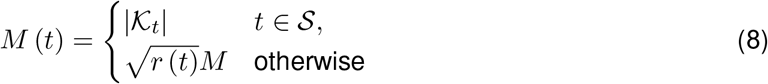

noting that *M* (*t*) ≤ ∑ _*t*′∈child(*t*)_ *M* (*t*′) due to the concavity of the square root. The concavity matters because it ensures that the size constraint becomes more restrictive as we move up the tree. In this case, the allowed budget increases faster than proportionally until 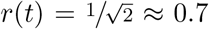 and slower than proportionally after. Thus, at lower ranks, many of the decisions for removing *k*-mers are left to its ancestors; note that a *k*-mer not removed at a taxon can still be removed further up. Close to the leaves, very few *k*-mers would be removed (e.g., a single species out of 10^4^ still is assigned 1% of the total budget when visiting this node), while close to the root, taxon budgets get closer to the full budget (e.g., a phyla with 2*/*3 of the species is assigned 82% of the budget). Also, going up the tree, the *k*-mer budget assigned to each taxon increases compared to each child, but it never exceeds the total budget assigned to its children. Clearly, *M* (root) matches the full budget allowed.

#### FilterByRank

For each hash taxon *t*, the FilterByRank function of Algorithm 1 removes *k*-mers when the size of the table **H**_*t*_ exceeds *M* (*t*). First, each row is pruned to *b k*-mers according to the strategy described below. If the table size is still greater than *M* (*t*), we continue removing *k*-mers, one at a time from each row in the descending order of their sizes, and iterating until the size constraint is satisfied.

For a *k*-mer set 𝒦, a taxonomy 𝒯, and its corresponding species set 𝒮, we define R, R′ : 𝒦 *×*𝒯 → [0, |𝒮|] as

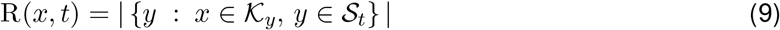

and

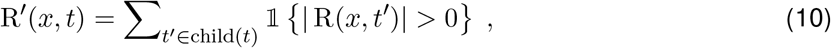

both illustrated in Fig. S2. We express each ranking strategy using either function and removing low-ranked *k*-mers accordingly. Discriminative and common *k*-mer strategies simply rank *k*-mers inversely and proportionally (respectively) to R or R′. In the case of ties, which can happen frequently, especially near leaves, we simply break them randomly.

The goal of covering all species can be imposed as a complex set covering problem. Instead, in the interest of scalability, we opt for a simple approach using weighted sums. We set

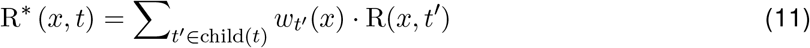

where *w*_*t′*_ down-weights groups that are highly sampled among surviving *k*-mers (see Fig. S2). A natural choice of weights is 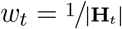. However, because |**H**_*t*_| is independent of the *k*-mer or the row being considered, this approach will consistently down-weight the children with the largest tables; in the extreme case, it can remove all *k*-mers from a child altogether. Thus, we define the weight of a *k*-mer *x* locally for its LSH table row by setting it to be inversely proportional to the *coverage* of each child taxon among *k*-mers of that row:

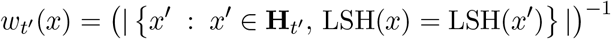

As a result, children covered with fewer *surviving k*-mers get higher weights, leading the process at the parent node to remove them less often (see Fig. S2 for a toy example). However, since decisions are made per row, and assignment of *k*-mers to rows is random, this avoids consistently removing the same species.

#### ExtractKmers

In extracting *k*-mers from all input genomes, we use minimizers to perform an initial subsampling by choosing the *k*-mer whose encoding has a MurmurHash3 (Appleby, 2009) value that is the smallest in a local window of size *w* = *k* + 3. We adopt the left/right encoding of *k*-mers introduced by Rachtman, Bafna, et al. (2021); this encoding allows fast calculation of Hamming distance using just four instructions (a pop-count, an XOR, an OR, and a shift). For *k >* 16, these encodings require 64 bits. However, *h* many *k*-mer positions are already given by the hash index and do not need to be saved as a part of the encoding. In KRANK, unlike CONSULT-II, we save memory by dropping these positions; this reduces the encoding size to a 32bit number when *k* − *h* ≥ 16, which is the case in our default configurations (*k* = 30 and *h* = 14 or *k* = 29 and *h* = 13). Since each *k*-mer encoding is only 32 bits, we save the encodings in the hash table; note that this is different from CONSULT-II, which saved pointers to a separate *encoding* array.

#### Populating LSH tables (FillTable, UnionTables, and ProjectTable)

For each leaf *t*, FillTable computes LSH and encoding values of each *k*-mer *x* ∈ *K*_*t*_ based on *h* positions, and stores *k*-mer’s compact encoding in the row indexed by the LSH value. We keep a counter initialized to 1 associated with each *k*-mer to enable computing R(*x, t*) later. For non-leaf taxa, UnionTables combines hash table rows by taking the union of *k*-mers. Luckily, the union operation can be defined on a per-row basis as the same *k*-mers share LSH value; independent union operations are performed in parallel across rows. Union also updates R(*x, t*). Furthermore, KRANK uses some of the available threads to perform post-order traversal in parallel.

Below root, KRANK saves rows of the hash tables using vectors that allow dynamic size changes during construction, switching to static structures only at the end. At the root, each row gets pruned to *b k*-mers, again using the ranking. This allows converting the table to a memory-efficient flattened static 1D array of size 2^2*h*^*b*.

#### Batching

Since we need to merge multiple tables at internal nodes, the required memory may exceed what is available. To tackle this issue, we used a batching approach. Except for the AdaptiveSizeCon-STRAINT, all components of our algorithm can be performed independently on a per-row basis. Therefore, we divide rows of the tables into batches and build them separately and in parallel. KRANK has a parameter to set the number of batches, and each batch’s size constraint is simply updated as *M* (*t*) divided by the number of batches.

### Datasets

#### WoL-v1 dataset (read classification)

We demonstrate the pros and cons of selection strategies using the microbial WoL-v1 dataset (Zhu et al., 2019) composed of 10,575 genomes (accessible at https://biocore.github.io/wol/download). We excluded 100 archaeal genomes used by Rachtman, Bafna, et al. (2021) as queries and five genomes with IDs missing from NCBI from the reference set, leaving us with 569 archaeal and 9901 bacterial references. We chose 756 queries in total; these include 666 bacterial genomes added to RefSeq after WoL-v1 was constructed, 10 randomly selected bacteria from the reference set, and all 80 of 100 archaeal queries from Rachtman, Bafna, et al. (2021) with Jaccard index above 0 to some reference genome. The queries span a range of distances to the closest reference genome and are binned into novelty groups as such (Fig. S1A). The query set covers 55 phyla and 396 genera, with some phyla highly represented (e.g., 253 from *Pseudomonadota*, 91 from *Bacillota*) but including several rare phyla (Fig. S1B). We generated 150bp reads using ART (Huang et al., 2012) at high coverage with default Illumina error profile and then subsampled down to 66667 reads for each query (≈ 2*×* coverage for these genomes). We use the NCBI taxonomy provided with the WoL-v1 dataset as our reference taxonomy.

We used this dataset to explore the various heuristics of KRANK and to also compare final KRANK versus other methods. We examined the same query set in both analyses. We compared KRANK against CONSULT-II v0.4.0 (Ş apcı et al., 2024), where *k*-mer selection is arbitrary, Kraken 2 (with default settings), and CLARK (Ounit, Wanamaker, et al., 2015) (with default settings, *k*=31; species rank), which uses discriminative *k*-mers. For KRANK, the classification algorithm is identical to CONSULT-II v 0.5.0. We tested KRANK with two memory levels: KRANK-hs (high-sensitivity) sets *k*=30, *h*=14, *b*=16 and uses 51.2Gb of memory while KRANK-lw (lightweight) sets *k*=29, *h*=13, *b*=16 and 12.8Gb of memory. For details, see the exact commands and tool versions in the supplementary information.

#### Profiling on CAMI-I high complexity dataset

This dataset contains five different high-complexity samples, each of size 75Gbp, which are simulated by mimicking the abundance distribution of the underlying microbial communities (Sczyrba et al., 2017). We tested the Bracken (Lu et al., 2017) extension of Kraken 2 which combines its results with Bayesian priors for better profiling. We used the same custom libraries built with the WoL-v1 dataset, with default parameters as described in WoL-v1 dataset (read classification). Results for CONSULT-II, CLARK, and Bracken are retrieved from our prior work and are available at https://github.com/bo1929/shared.CONSULT-II.

#### Profiling on CAMI-II marine and strain-madness datasets

We used the ten-sample marine dataset (5Gbp each, available at https://frl.publisso.de/data/frl:6425521/marine/) and the 100-sample (2Gbp each, available at https://frl.publisso.de/data/frl:6425521/strain/) strain-madness dataset, which are the two main datasets focused on abundance profiling in the CAMI-II challenge. For all tools except CONSULT-II and KRANK, we used CAMI-II results submitted to the challenge available at https://github.com/CAMI-challenge/second_challenge_evaluation. We exclude versions of the methods that were submitted to the first challenge.CONSULT-II results were again retrieved from https://github.com/bo1929/shared. CONSULT-II. We built and used a KRANK library in high-sensitivity setting (*k*=30, *h*=14, *b*=16) using 72,766 genomes selected from NCBI RefSeq snapshot as of 2019/01/08 (were retrieved from https://openstack.cebitec.uni-bielefeld.de:8080/swift/v1/CAMI_2_DATABASES/) by allowing each species to contribute at most 500 genomes (otherwise randomly selected among available genomes).

#### Evaluation metrics

We describe evaluation metrics, referring the reader to Supplementary Material for further commands and details.

#### Read classification

We evaluated the read classification with respect to the NCBI taxonomy. Each prediction is separately evaluated across each taxonomic rank. When the reference set had at least one genome matching the query taxon at a particular rank, we call it a *positive*: *TP* if it is the correct taxon, *FP* if it is the incorrect taxon, and *FN* if the read is not classified at that rank. Similarly, at a particular rank, when the reference set lacks a genome from the query taxon, we call it a *negative*: *TN* if the read is not classified at that rank, *FP* if it is classified, which would necessarily be false. We ignored queries at ranks where the true taxonomic ID given by NCBI is a missing rank.

We use custom scripts to process the specific output format of each tool and compare it with the taxonomy used to build reference libraries. These scripts are available on https://github.com/bo1929/shared.KRANK. We also provide distances of each query genome to the closest reference genome *d*\* as estimated by Mash (Ondov et al., 2016), taxonomy files, and the outputs of evaluation scripts.

#### Taxonomic profiling

To measure taxonomic profiling performances on both CAMII challenges, we reported the same metrics as those emphasized in original publications. We used OPAL (version 1.0.12) to compute the metrics with -n option (which renormalizes after removing the unclassified portion).

On CAMI-I, we used the main two metrics singled out by the open-community profiling assessment tool (OPAL, available on GitHub: https://github.com/CAMI-challenge/OPAL) (Meyer, Bremges, et al., 2019): the Bray-Curtis dissimilarity between the estimated profile and the true profile and Shannon’s equitability as a measure of alpha diversity. All the results are averaged across all 5 samples. On the CAMI-II Challenge, we followed the original paper by Meyer, Fritz, et al. (2022) and compared methods based on L1 norm error and weighted UniFrac error. Both L1 norm error and weighted UniFrac error measure the accuracy of the relative abundance estimations of taxa in a sample. The UniFrac metric is computed across the entire taxonomic tree, while the L1 norm is computed at each taxonomic rank. The weighted UniFrac has a maximum of 16 on this dataset, so we report 16 - UniFrac; similarly, the L1 norm can never be more than 2, so we report 2 - L1. We report averages across 10 samples and 100 samples respectively for marine and strain-madness datasets.

## DATA ACCESS

Simulated query sequences for read classification can be found at https://ter-trees.ucsd.edu/data/krank/KRANK-queries.tar.gz. Our results, metadata files and scripts used to create figures, and custom scripts used to evaluate read classification results are available at https://github.com/bo1929/shared.KRANK. KRANK’s software can be found on GitHub: https://github.com/bo1929/ KRANK. We also share both the software and custom auxiliary scripts as Supplementary Code. The list of KRANK libraries used across our experiments can be found at https://ter-trees.ucsd.edu/data/krank/, a tutorial for running KRANK, and descriptions of these libraries are available on GitHub.

