## Supplementary Information for "Memory-bound *k*-mer selection for large and evolutionary diverse reference libraries"

### Supplementary Methods

#### Evaluation metrics and tools used

To compute recall, precision, and F1 for read classification, we used the following commands, which can be run after setting the current working directory to `https://github.com/bo1929/shared.KRANK`:

1. `python scripts/evaluate.py --output-dir $OUTPUT_DIR \`
   `--taxonomy-database-path data/ReferenceTaxonomy-nodes.dmp --tvth 0.03 \`
   `--reference-ranks-path data/10kBacteria-ranks_tid.tsv \`
   `--query-ranks-path data/uDance-ranks_tid.tsv --results-path $RESULTS_FILE`
   where `$RESULTS_FILE` is the output report of CONSULT-II, including the prediction for each read, and `$OUTPUT_DIR` is simply the output directory,
2. `python scripts/summarize_evaluations.py $OUTPUT_FILE \`
   `data/dist-to-closest.txt 1 > $EVALUATION_SUMMARY`
   where `$OUTPUT_FILE` is the output of the previous command, and `$EVALUATION_SUMMARY` is the summary results across all reads used to generate figures.

For the CAMI-II challenge, we ran the commands given at `https://github.com/CAMI-challenge/second_challenge_evaluation/tree/master/profiling`, using the results of all participants provided in the same repository. For CAMI-I, we used the ground-truth given at `https://github.com/CAMI-challenge/OPAL/blob/master/data/gs_cami_i_hc.profile`. To post-process abundance estimates of each tool to the CAMI profiling Bioboxes format, we used scripts found at `https://github.com/CAMI-challenge/docker_profiling_tools`.

#### Parameter configurations of tools

For CONSULT-II, we used $l$=2 and $k$=32 (minimized from canonical 35-mers) for all memory levels. We set hash table parameters $h$ and $b$ to `-h 15 -b 7`, `-h 13 -b 14`, `-h 12 -b 16`, and `-h 12 -b 9`, respectively for 140.7Gb, 16Gb, 5Gb and 3Gb libraries. For other parameters (LCA probability function and maximum match distance), default values were used.

As Rachtman, Balaban, et al. (2020) found default settings to be preferable for query identification, we used default settings for Kraken 2 (`--kmer-len 35`, `--minimizer-len 31`, and `--minimizer-spaces 7`) and CLARK ($k$=31, default classification mode, species rank for classification, and `-m 1`). We used `--max-db-size` parameter to adjust Kraken 2 database sizes (4Gb, 16Gb, and default 46Gb).

For KRANK, we used the following parameter configurations for corresponding library sizes; 76.8Gb: `-k 30 -w 33 -h 14 -b 16` with $l$=3, 51.2Gb: `-k 30 -w 33 -h 14 -b 16` with $l$=2 (high-sensitivity), 25.6Gb: `-k 30 -w 33 -h 14 -b 16` with $l$=1, 12.8Gb: `-k 29 -w 32 -h 13 -b 16` with $l$=2 (lightweight), 6.4Gb: `-k 29 -w 32 -h 13 -b 16` with $l$=1, 3.2Gb: `-k 28 -w 31 -h 12 -b 16` with $l$=2, 1.6Gb: `-k 28 -w 31 -h 12 -b 16` with $l$=1.

#### Software versions and commands used

Here, we provide the exact commands that we used to run external tools, CONSULT-II, and KRANK throughout our experiments, together with their version information.

##### Genomic distance estimation using Mash (Ondov et al., 2016)

We used Mash (version `2.3`) to estimate genomic distances. To create a Mash sketch from a genome and then use it to estimate genomic distance, we used the below commands.

1. `mash sketch -k 30 -s 750000 -p $NTHREADS -o $SKETCH_FILE $INPUT_FASTA`

2. `mash dist $SKETCH_FILE1 $SKETCH_FILE2`

**Short read simulation with reference using ART (Huang et al., 2012)**

We simulated short reads with length $L$=150 and coverage $c$, with the default error and quality profiles of Illumina HiSeq 2500 using ART (version 2.5.8 – single read mode) with the command below.

```
art_illumina -ss HS25 -l 150 -f c -na -s 10 -i $INPUT_FASTA -o $OUTPUT_FASTQ
```

**Extraction of $k$-mer sets using Jellyfish (Marçais and Kingsford, 2011)**

We used Jellyfish (v2.3.0) to compute distinct $k$-mer sets and $k$-mer profiles (for canonical 32-mers and 35-mers) of concatenated reference genomes (in FASTA format). The command is given below.

```
jellyfish count -m 35 -s 100M -t 24 -C $INPUT_FASTA -o $COUNT_FILE
```

Then, to output the list of distinct $k$-mers together with their counts into a FASTA file, we used the following command. Note that we do not use count values.

```
jellyfish dump $COUNTS_FILE > $OUTPUT_FASTA
```

**Downsampling reads with `seqtk` (Li, 2018)**

To subsample read collection down to a specified number of reads, denoted by $n$ here, we used `seqtk` (version 1.3r106) with the command below.

```
seqtk sample -s150 INPUT_FASTQ n > OUTPUT_FASTQ
```

**Building a KRANK library and querying sequences**

The version of KRANK we presented in this paper is v0.4.0. Running the following commands in order creates a KRANK library that can be queried against to find $k$-mer matches.

1. We first initialize a library, and preprocess all input genomes to extract $k$-mers. We built two libraries to use together: one with `--adaptive-size` option and the other with `--free-size` option, random seeds set to 0 and 19, respectively. We set $k$-mer length (`-k`) to 30, minimizer window size (`-w`) to 33, number of LSH positions (`-h`) to 14, number of table columns (`-b`) to 16, and number of bits to determine batch IDs (`-s`) to 9 (hence $2^9$ batches in total). Two files are required to be given with arguments (`-t` and `-i`): `$TAXONOMY_NODES` is an NCBI taxonomy `nodes.dmp` file and `$INPUT_MAP_FILE` is a tab-separated file, its first column is for the species IDs and the second column is for the genome paths. The option `--input-sequences` specifies that input files are genomes, not $k$-mers extracted by some external tool, `--from-scratch` specifies that the library is to be initialized.
```
krank --seed $RANDOM_SEED build -k 30 -w 33 -h 14 -b 16 -s 9 \
--l $LIBRARY_DIR -t $TAXONOMY_NODES -i $INPUT_MAP_FILE \
--kmer-ranking 1 --adaptive-size \
--from-scratch --input-sequences --num-threads $NTHREADS
```
2. Once the library is initialized, each batch can be built separately (i.e., `--target-batch` 1 to 512) using the below command. Note that this command also adds the soft-LCA information to the library.
```
krank --seed $RANDOM_SEED build \
--l $LIBRARY_DIR -t $TAXONOMY_NODES -i $INPUT_MAP_FILE \
--kmer-ranking 1 --adaptive-size -from-library --num-threads $NTHREADS \
--target-batch $BATCH_NUM --input-sequences
```
3. Lastly, we queried sequences against the constructed library and generated a CONSULT-II compatible $k$-mer match information file for further classification and profiling.
```
krank query -l $LIBRARY_DIR -o $OUTPUT_DIR -q $QUERY_PATHS_FILE \
--save-match-info --max-match-distance 5 --num-threads $NTHREADS
```

**Taxonomic identification with Kraken 2 (Wood et al., 2019) and Bracken (Lu et al., 2017)**

We used `v2.1.3` and `v2.8` for Kraken 2 and Bracken in our experiments with their default parameters, respectively. For CAMI-II (Meyer, Bremges, et al., 2019), the presented results are for Bracken `v2.2`, which was the version submitted to the challenge.

In order to construct a custom Kraken 2 reference library from WoL-v1 genomes, and to extend it with Bracken for abundance profiling, we used the following commands:

1. We first initialized a library named `$DBNAME`.
   ```
   kraken2-build --download-taxonomy --db $DBNAME
   ```
2. Next, we changed the downloaded taxonomy to match the taxonomy files provided with the WoL-v1 database, retrieved from https://biocore.github.io/wol/data/taxonomy/.
3. We then added our custom genomes to the reference database.
   ```
   find $REFERENCE_GENOMES_DIR -name '*.fna' -print0 | \
   xargs -0 -I -t -n1 kraken2/kraken2-build --add-to-library  --db $DBNAME
   ```
4. After that, we built the database with specified $k$-mer & minimizer lengths `$KMER_LEN` (35) & `$MINIMIZER_LEN` (31) and default number of wind-carding positions $s = 7$.
   ```
   kraken2-build --build --db $DBNAME --threads 14 --max-db-size $MAX_SIZE \
   --kmer-len $KMER_LEN --minimizer-len $MINIMIZER_LEN --minimizer-spaces s
   ```
5. Lastly, given read length `$READ_LEN` (150), we processed the constructed library with Bracken.
   ```
   ./bracken-build -d $DBNAME -t $NTHREADS -k $KMER_LEN -l $READ_LEN
   ```

To query against the Kraken 2 reference library for read classification and Bracken for abundance profiling, we used the commands below.

1. ```
   kraken2 --use-names --threads $NTHREADS --report $REPORT_FILE \
   --db $DBNAME $QUERY_FASTQ > $KRAKEN_OUTPUT
   ```
2. ```
   bracken -d $DBNAME -i $KRAKEN_OUTPUT -o $BRACKEN_OUTPUT \
   -r $READ_LEN -l S -t $NTHREADS
   ```

**Taxonomic identification with CLARK (Ounit, Wanamaker, et al., 2015)**

We used CLARK `v1.2.6.1` with its default parameters. CLARK failed to set target IDs for 494 genomes out of 10,470 WoL-v1 genomes during the construction of its database. These genomes were added manually by setting their corresponding taxon.

We followed the below steps to build a custom CLARK database in the directory `$DIRDB`.

1. Create the directory `Custom/` inside `$DIRDB` directory.
2. Copy reference WoL-v1 genomes with accession numbers into the `Custom/` directory.
3. Similar to Kraken 2, we set the taxonomy to the one provided with the WoL-v1 database, retrieved from https://biocore.github.io/wol/data/taxonomy/.
4. We ran `./set_target.ssh $DIRDB custom --species`.

We queried sequences against the CLARK database using the default mode (i.e., `-m 1`), and then used these classification results to estimate abundances by running the commands below.

1. `./classify_metagenome.sh -O $QUERY_FASTQ -R $RESULT_FILE -n $NTHREADS -m 1`
2. `./estimate_abundance.sh -F $RESULT_FILE -D $DIRDB`

**Taxonomic identification with CONSULT-II (Şapcı et al., 2024)**

We used CONSULT-II version `v0.4.0` for all experiments except profiling with KRANK databases, for which we used version `v0.5.0`. Note that, `v0.5.0` was not used in the original manuscript of CONSULT-II and differs from `v0.4.0` mainly at species-level abundance profiling. For CAMI-II profiling, we directly used results presented in Şapcı et al. (2024) (for CONSULT-II) and Meyer, Fritz, et al. (2022) (for all other tools).

We used the commands below to build CONSULT-II reference libraries.

1. First, we minimized canonical reference 35-mers (output of `jellfish dump`) to 32-mer minimizers with a custom script (version `v0.3.0`), provided with CONSULT-II.
   ```
   minimize -i $INPUT_35MERS_FASTA -o $OUTPUT_32MERS_FASTA
   ```
2. For the default 140.7Gb memory level, we used the command below to create the library. Here, `$INPUT_FASTA` is the output of above command: `$OUTPUT_32MERS_FASTA`. For 16Gb and 5Gb versions, one can set `-h 13 -b 14` and `-h 12 -b 16`, respectively.
   ```
   consult_map --input-fasta-file $INPUT_FASTA --output-library-dir $LIBDIR \
   --number-of-positions 15 -t 2 -l 2 -b 7.
   ```
3. After constructing the reference library, we added taxonomic information (soft-LCA information per $k$-mer) by using the following three commands:
   (a) Adding soft-LCAs requires an auxiliary file (`$TAXONOMY_LOOKUP`) that can be generated from a taxonomy file (`$TAXONOMY_NODES`, i.e., `nodes.dmp`), using a script CONSULT-II provides.
   ```
   python scripts/construct_taxonomy_lookup.py \
   -from-taxonomy-database $TAXONOMY_NODES
   ```
   (b) Then, running these two commands in order would finalize the library construction. Note that `$REFERENCE_GENOMES_DIR` is the directory in which all reference genomes are stored as FASTQ files.
   ```
   consult_search -i $LIBDIR --taxonomy-lookup-path $TAXONOMY_LOOKUP \
   -q $REFERENCE_GENOMES_DIR --filename-map-path $FILENAME_MAP \
   --thread-count $NTHREADS --init-ID
   consult_search -i $LIBDIR --taxonomy-lookup-path $TAXONOMY_LOOKUP \
   -q $REFERENCE_GENOMES_DIR --filename-map-path $FILENAME_MAP \
   --thread-count $NTHREADS --update-ID
   ```

Performing taxonomic classification and profiling contains two stages: finding $k$-mer matches and reporting them in an appropriate file format and then utilizing them to predict read-level assignments or to estimate sample-level composition estimates.

1. The below command outputs the match information files per query (`$MATCH_INFO`), with filenames that are query filenames prefixed with `match-info_*`, to `$OUTPUT_DIR`. This file format can also be generated by KRANK, and hence, the next steps are identical to CONSULT-II.
   ```
   consult_search -i $LIBDIR -q $QUERY_FASTQ -o $OUTPUT_DIR \
   --maximum-distance 5 --save-matches --thread-count $NTHREADS
   ```
2. To classify each read, simply run:
   ```
   consult_classify -i $MATCH_INFO -o $OUTPUT_DIR \
   --taxonomy-lookup-path $TAXONOMY_LOOKUP
   ```
3. For profiling, the below two commands are needed.
   (a) We first predict the number of reads belonging to each taxon.
   ```
   consult_profile -i $MATCH_INFO -o $OUTPUT_DIR \
   --taxonomy-lookup-path $TAXONOMY_LOOKUP
   ```
   (b) Then, using the output of the above command, we perform normalization and genome size correction. Here, `$THRESHOLD` (1000 in our case) is the total vote threshold for a taxon to be included, `$ABUNDANCE_REPORT` is the `consult_profile`'s output, `$NCBI_TAXONOMY_DIR` is a directory which stores `nodes.dmp` and `names.dmp` files, `$METADATA_FILE` is a tab seperated file which is required to contain `species_id` and `total_length` columns for each reference genome. The `total_length` corresponds to the genome size of the genome belonging to species `species_id`. `$S_ID` is a sample identifier name for the header of the report.
   ```
   python scripts/postprocess_profiles.py $THRESHOLD $ABUNDANCE_REPORT \
   $NCBI_TAXONOMY_DIR $METADATA_FILE $S_ID
   ```

### Supplementary Figures

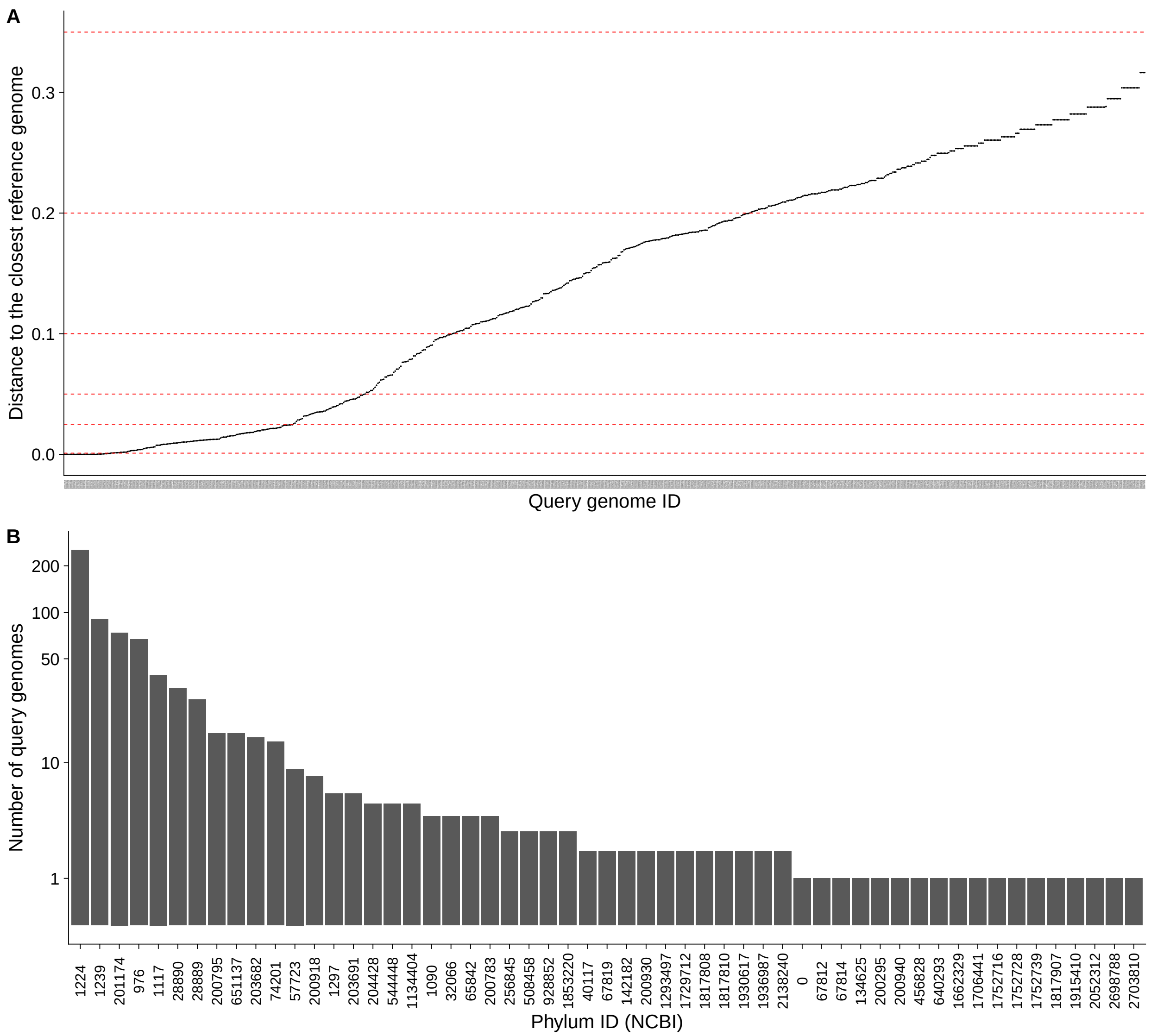


Figure S1: A) The distance to the closest reference genome $d^*$, as estimated by Mash, of each query genome. Each data point is for a query genome ($x$-axis). Dotted horizontal lines show boundaries of bins. B) Distribution of 765 queries across different phyla. The number of query genomes from the corresponding phylum is given the $y$-axis, and the $x$-axis is the NCBI ID of the phylum. 0 refers to genomes with unknown/unassigned groups at the phylum level. The $y$-axis is in one plus log scale.

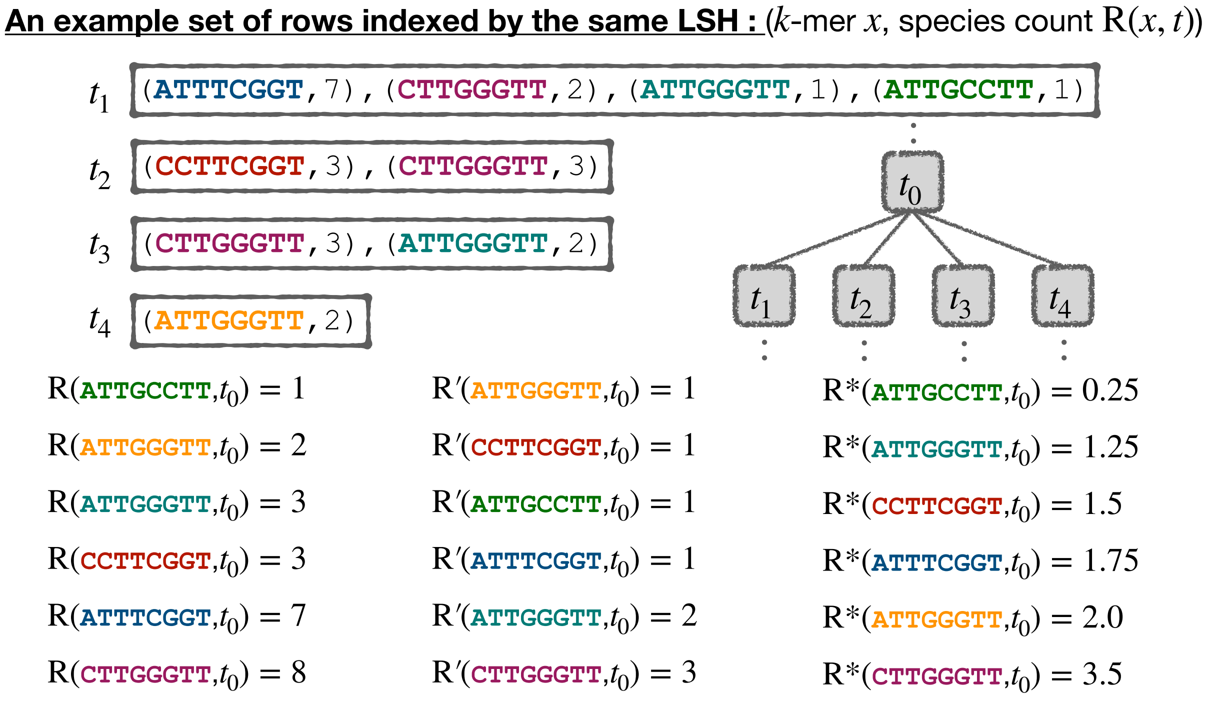


Figure S2: An illustration of a toy example to compare different $k$-mer ranking strategies: $\mathrm{R}$, $\mathrm{R}'$, and $\mathrm{R}^*$. Tables of sibling taxa $t_1$, $t_2$, $t_3$, and $t_4$ are being merged for parent $t_0$'s table. For a particular LSH index, $t_1$, $t_2$, $t_3$, and $t_4$ have 4, 2, 2, and 1 $k$-mers in their corresponding row, respectively. Hence, weights for $\mathrm{R}^*$ will be $^1/_4$, $^1/_2$, $^1/_2$, and $1$.

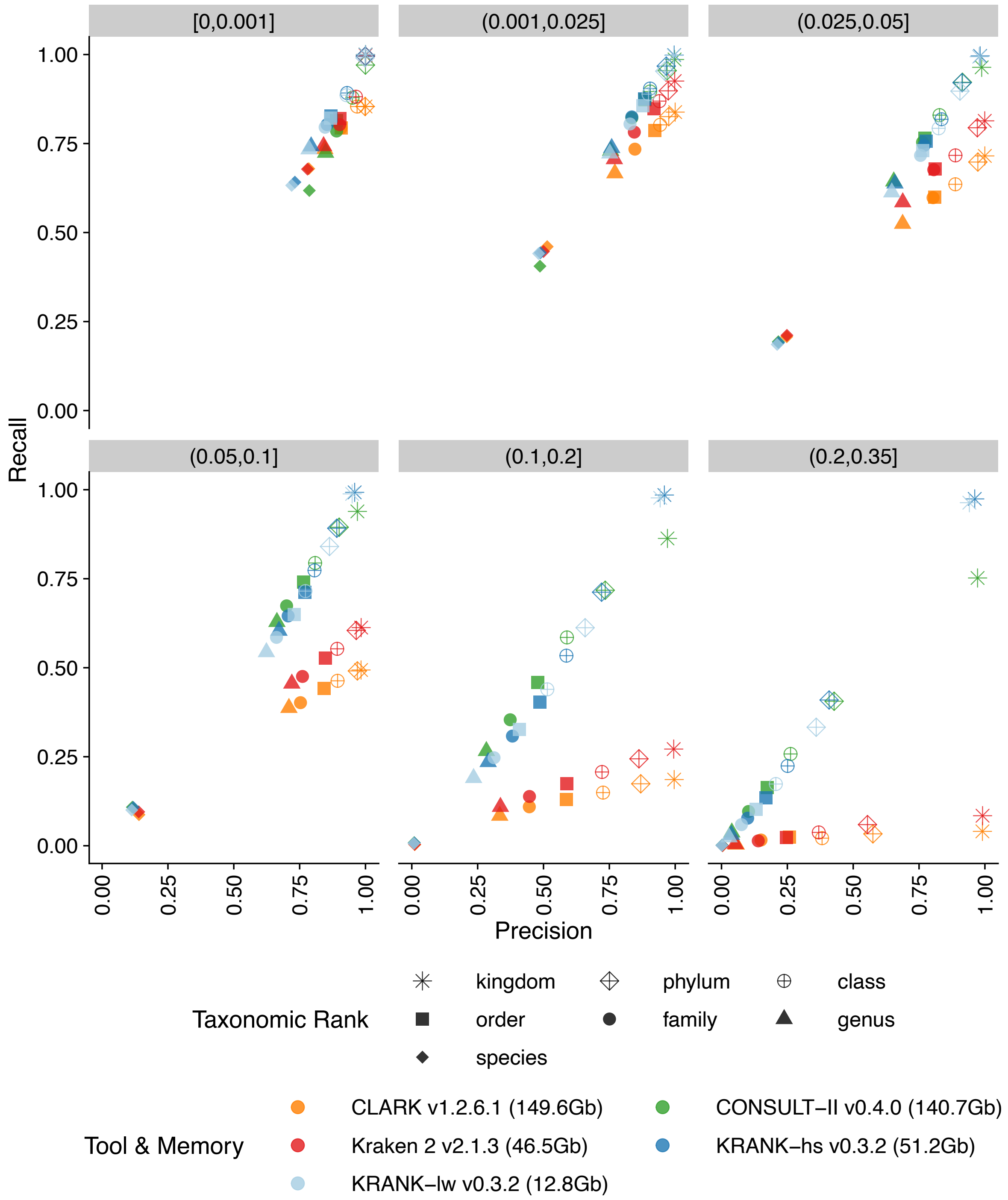


Figure S3: A more fine-grained comparison of the high-sensitivity (hs) and the lightweight (lw) modes of KRANK with other methods across different ranks on the WoL-v1 dataset with 66667 simulated 150bp reads from each of 756 query genomes. Results are averaged over all queries and reported per novelty bins ($d^*$; see Fig. S1) and ranks in terms of precision ($x$-axis) and recall ($y$-axis) scores.

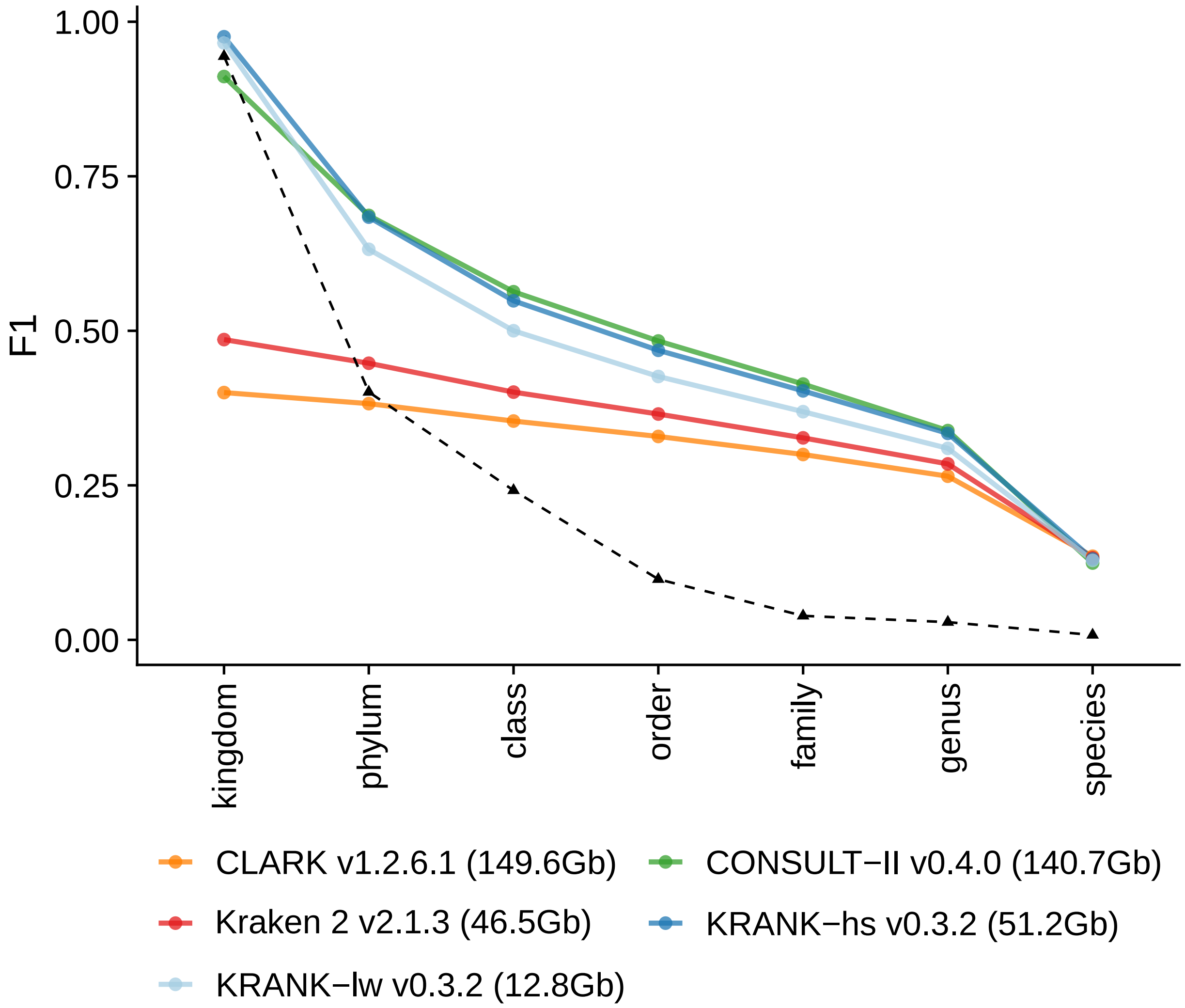


Figure S4: Comparison of KRANK with other methods across different ranks on the WoL-v1 dataset with 66667 simulated 150bp reads from each of 756 query genomes. Results are averaged over all queries. The dashed line stands for the *null model* which assigns all sequences to the largest taxon (in terms of number of query genomes) in the query set for the corresponding rank.

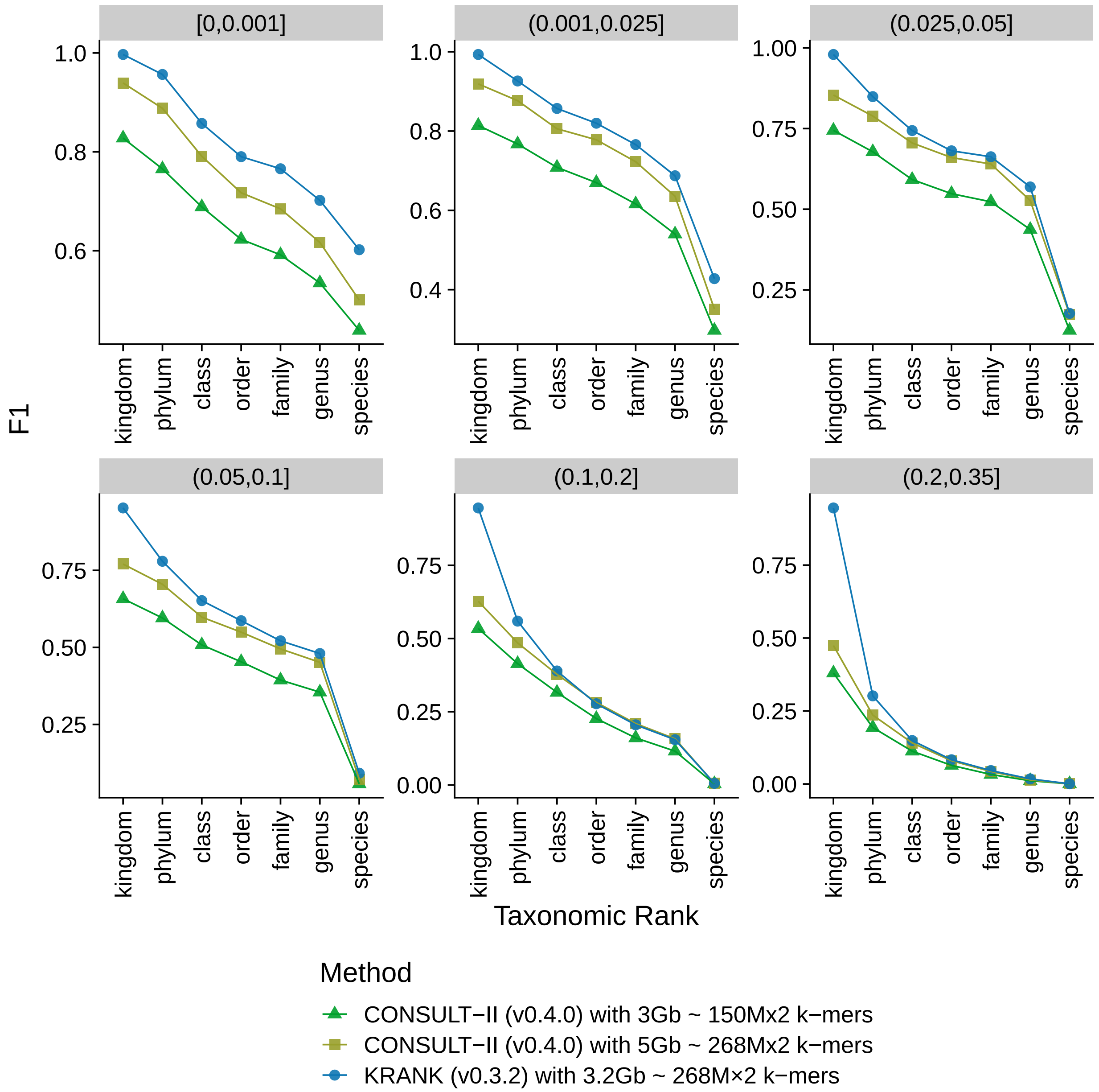


Figure S5: Comparison of CONSULT-II and KRANK in cases with the same memory level and the same number of $k$-mers in the library. KRANK can store 268M $k$-mer in a single table using less memory (1.8Gb), but the selected $k$-mer set further improves the performance regardless of the memory. F1 scores are reported across all ranks as averages of all queries in each novelty bin.

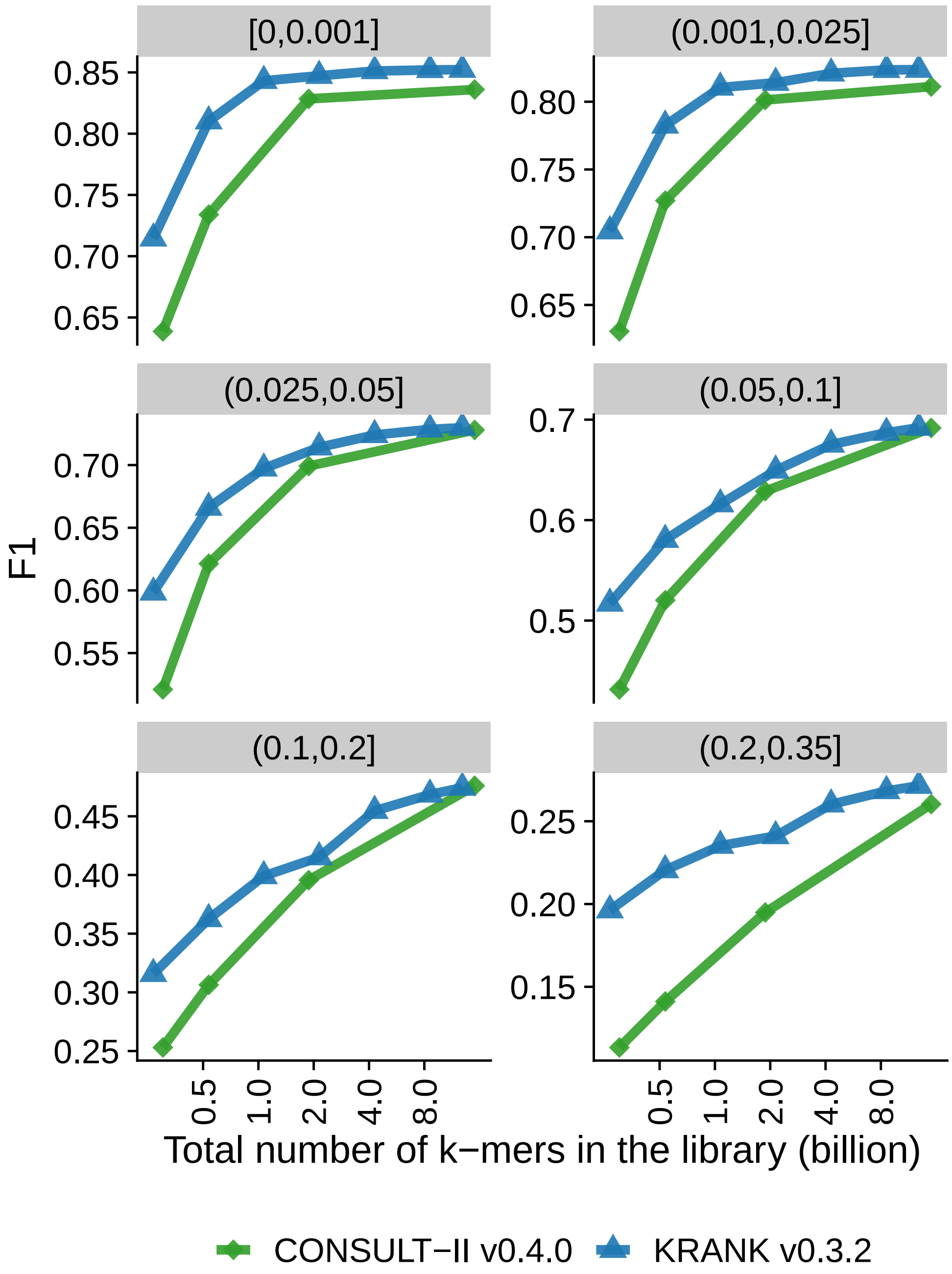


Figure S6: Comparison of classification performances CONSULT-II and KRANK as the total number of non-distinct $k$-mers kept in the library. Each panel is a novelty bin, corresponding to the minimum distance to any reference genome $(d^*)$, measured by Mash (Ondov et al., 2016). F1 scores are reported across all ranks as averages of all queries in each novelty bin.

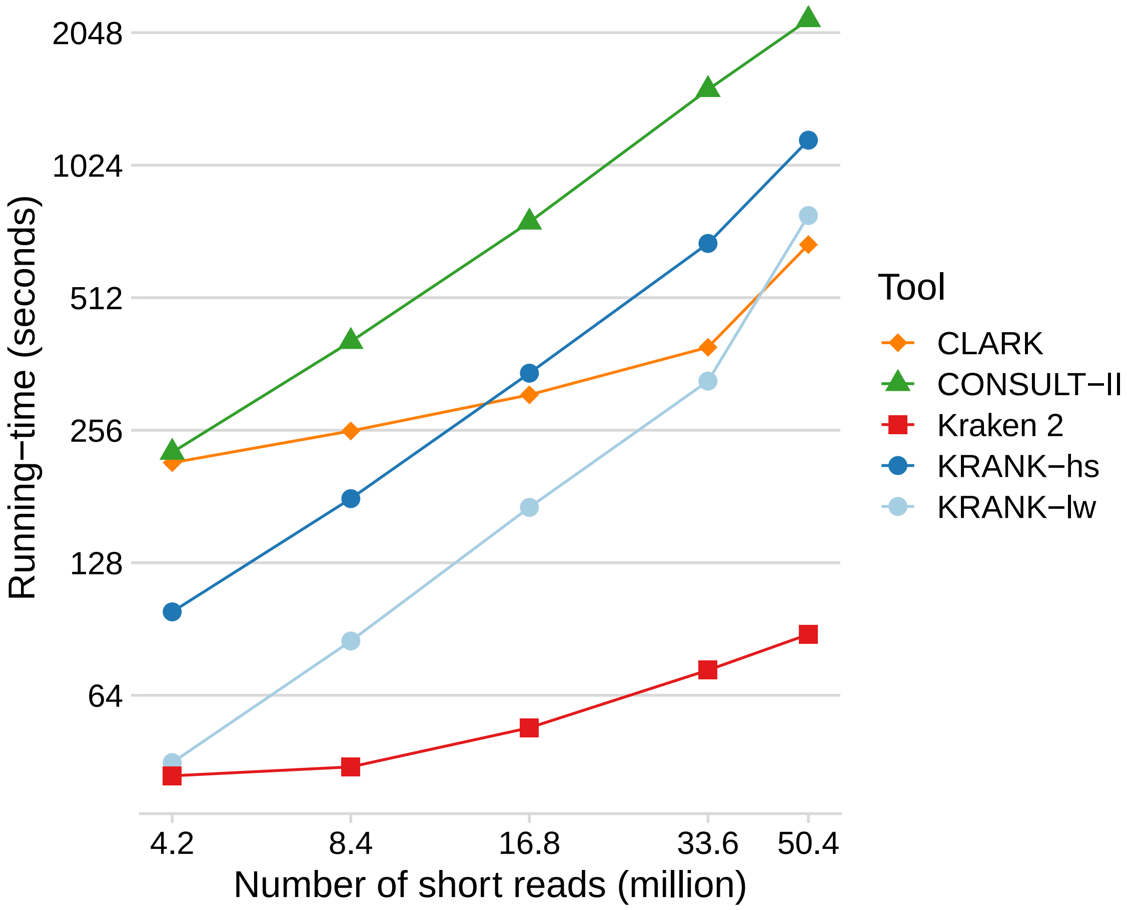


Figure S7: Query time for varying numbers of short reads sampled across 756 genomes from different novelty levels. Running time measurements were performed using 96 threads on a machine with 2.2 GHz AMD EPYC 7742 processors. Both axes ($x$ and $y$) are in log-scale.

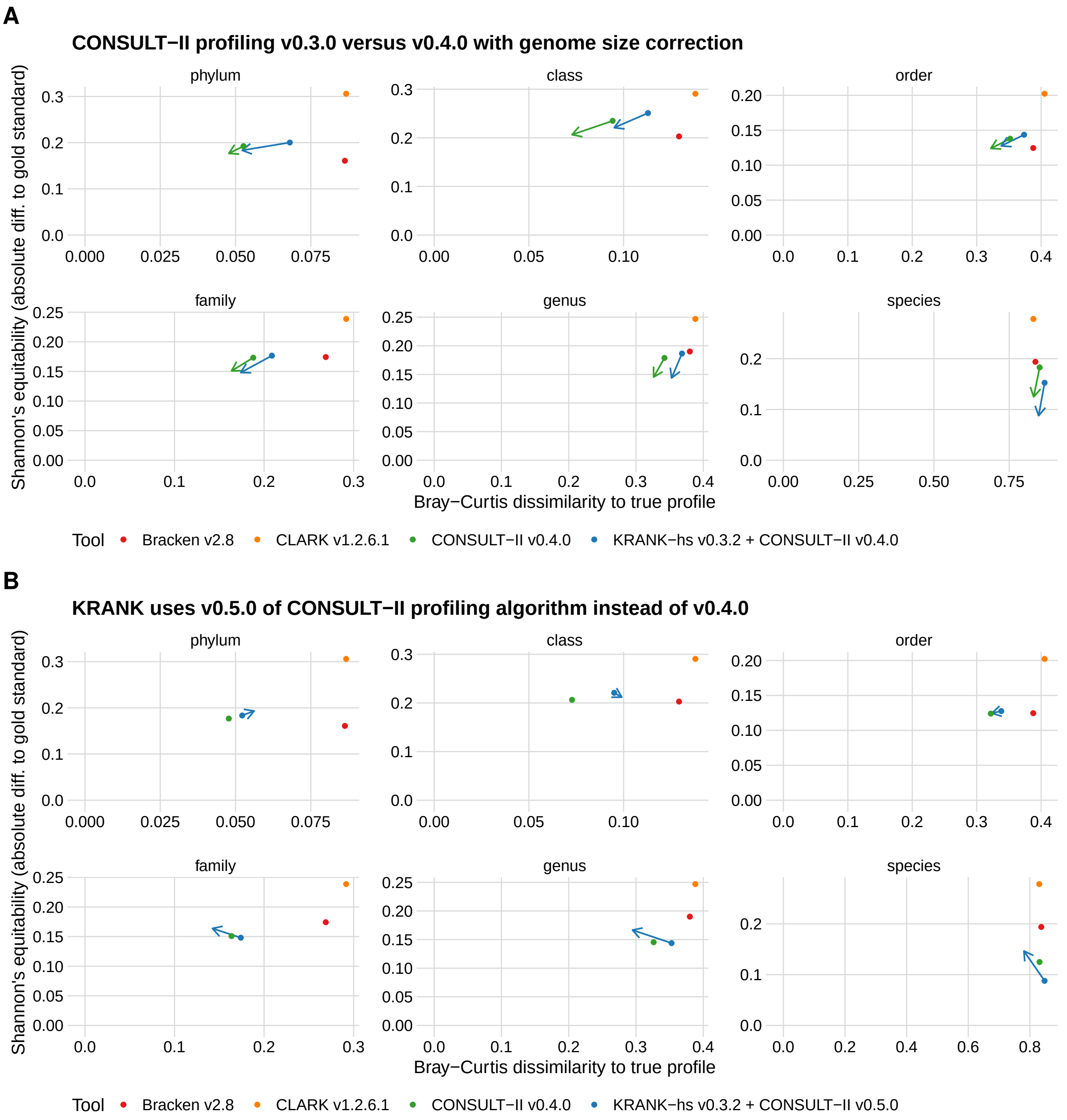


Figure S8: CONSULT-II and KRANK improvements across versions on CAMI-I dataset, based on Bray-Curtis dissimilarity to the true profile and Shannon's equitability difference with the gold standard. The bottom-left corner is perfect accuracy. A) CONSULT-II v0.4.0 versus v.0.3.0. The arrows show v0.3.0 (dot) and v0.4.0 (arrowhead) performance for KRANK and CONSULT-II. Applying genome size correction as a post-processing step in v0.4.0 improves v0.3.0. Note that, we use v0.4.0 for CONSULT-II profiling results, and v0.5.0 for KRANK profiling results. B) KRANK performance when using CONSULT-II v0.4.0 versus v0.5.0 for the profiling step. We use v0.5.0 and 0.4.0 throughout the manuscript respectively for KRANK and CONSULT-II profiling. We use v.0.4.0 for CONSULT-II because this version is used by Şapcı et al. (2024) and improvements of v0.5.0 were developed as part of this paper.

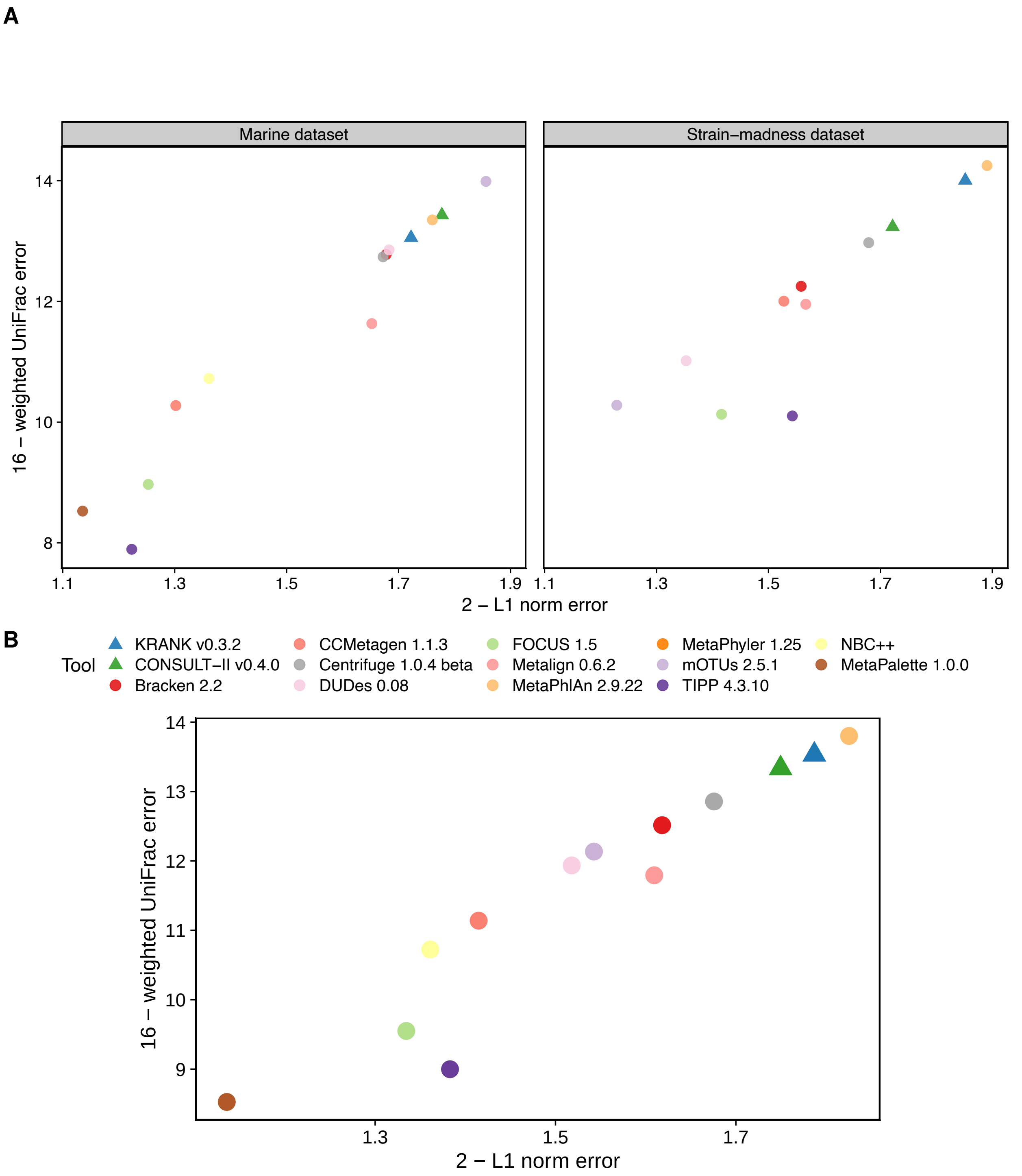


Figure S9: Comparing KRANK (high-sensitivity setting using 51.2Gb) with other participants in CAMI-II benchmarking challenge (Meyer, Fritz, et al., 2022). We report average L1 norm error averaged across all ranks and the rank invariant weighted UniFrac error. A) Each data point stands for the average of 100 strain-madness samples or 10 marine samples. B) Each data point stands for the average of two datasets: the average of 100 strain-madness samples and the average of 10 marine samples. Metrics were computed using OPAL with default settings and the `-n` option.

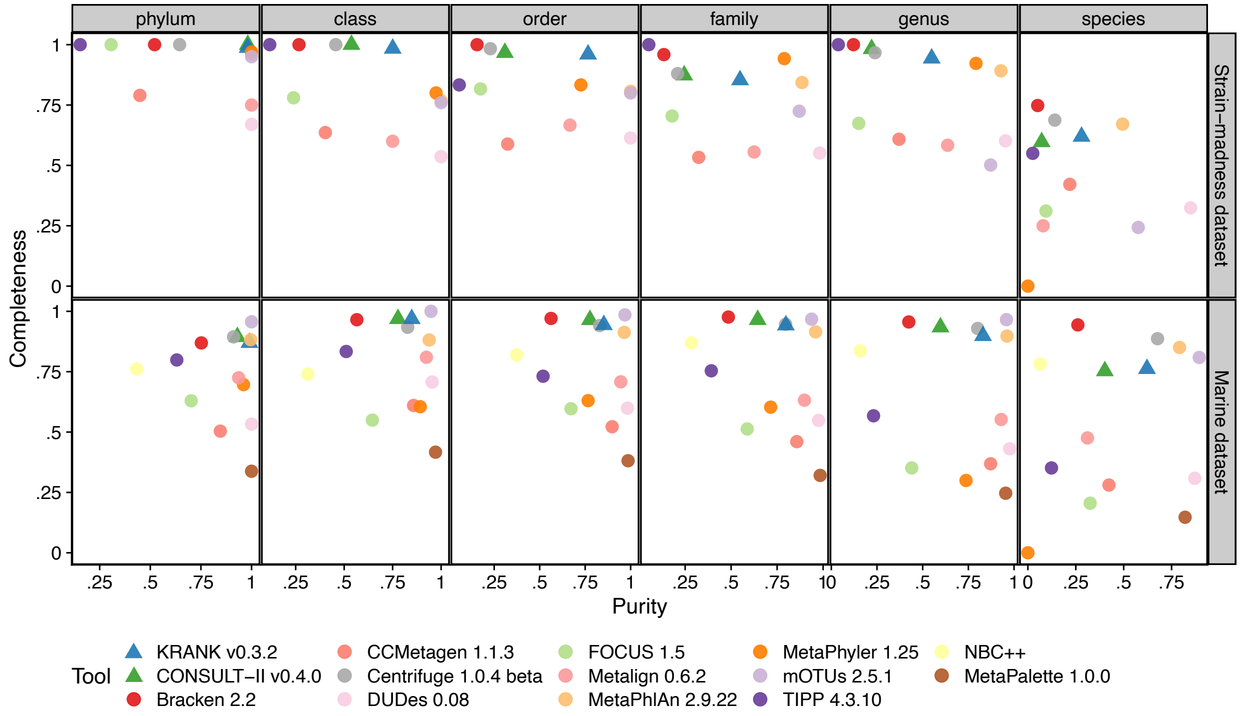


Figure S10: Comparing KRANK (high-sensitivity setting using 51.2Gb) with other participants in CAMI-II benchmarking challenge (Meyer, Fritz, et al., 2022). As in the original manuscript, we show purity versus completeness. Each data point stands for the average of 100 strain-madness samples or 10 marine samples. Metrics were computed using OPAL with default settings and the `-n` option.

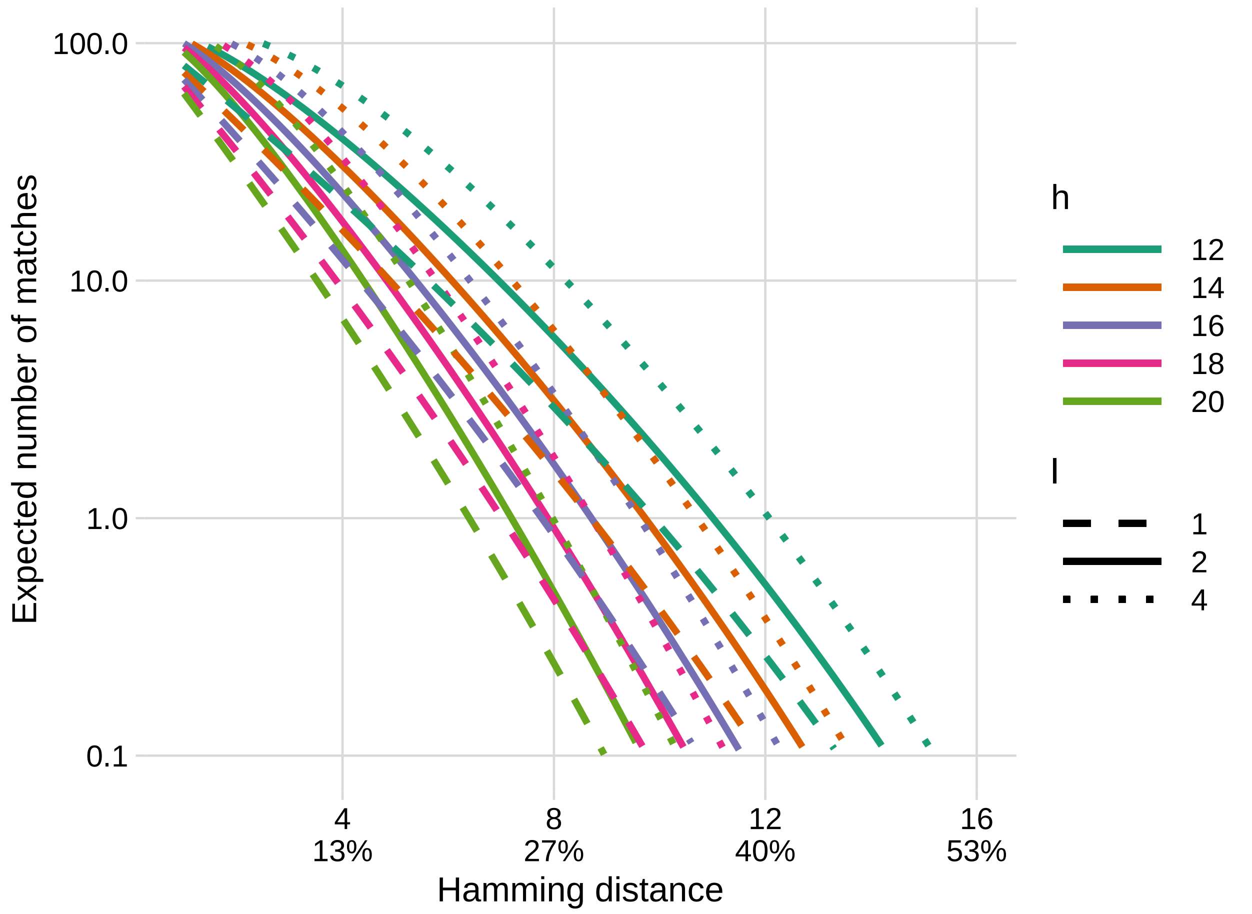


Figure S11: The expected number of 30-mers matched by the LSH approach for a short read of length $L{=}150$ as the normalized distance $^d/_k$ of the read to the closest match varies: $(L - k + 1)\rho(d)$. Lines show different settings of $l$ and $h$ for an infinite $b$, i.e., all reference $k$-mers are stored in the library.
